# Selective Microautophagy of Proteasomes is Initiated by ESCRT-0 and Is Promoted by Proteasome Ubiquitylation

**DOI:** 10.1101/2021.09.21.461272

**Authors:** Jianhui Li, Mark Hochstrasser

## Abstract

The proteasome is central to proteolysis by the ubiquitin-proteasome system under normal growth conditions but is itself degraded through macroautophagy under nutrient stress. A recently described AMPK (AMP-activated protein kinase)-regulated ESCRT (endosomal sorting complex required for transport)-dependent microautophagy pathway also regulates proteasome trafficking and degradation in low glucose conditions in yeast. Aberrant proteasomes are more prone to microautophagy, suggesting the ESCRT system fine-tunes proteasome quality control under low glucose stress. Here we uncover additional features of the selective microautophagy of proteasomes. Genetic or pharmacological induction of aberrant proteasomes is associated with increased mono- or oligo-ubiquitylation of proteasome components, which appear to be recognized by ESCRT-0. AMPK controls this pathway in part by regulating the trafficking of ESCRT-0 to the vacuole surface, which also leads to degradation of the Vps27 subunit of ESCRT-0. The Rsp5 ubiquitin ligase contributes to proteasome subunit ubiquitylation, and multiple ubiquitin-binding elements in Vps27 are involved in their recognition. We propose that ESCRT-0 at the vacuole surface recognizes ubiquitylated proteasomes and initiates their microautophagic elimination during glucose depletion.

**Saummary statement:** ESCRT-0 selectively targets aberrant proteasomes for microautophagy by recognition of proteasome ubiquitylation status to fine-tune proteasome quality control under low glucose conditions.

## Introduction

The ubiquitin-proteasome system (UPS) is a highly conserved proteolytic system that selectively degrades cellular proteins (Lecker et al., 2006). This pathway involves a cascade of reversible enzymatic reactions that covalently conjugates ubiquitin to substrates and targets them to the proteasome for degradation. Deubiquitylating enzymes (DUBs) remove ubiquitin from the substrates for recycling before the substrates are degraded by the proteasome into small peptides (Tomko and Hochstrasser, 2013). The 26S proteasome is composed of the 20S core particle (CP) and the 19S regulatory particle (RP), and the RP in turn comprises base and lid subcomplexes (Budenholzer et al., 2017). Dysfunction of the UPS has been implicated in the pathogenesis of many human diseases, including various cancers and neurodegenerative disorders (Thibaudeau and Smith, 2019).

Proteasomes are dynamic structures that change when cells are in distinct nutrient environments. Proteasomes are concentrated in the nucleus in yeast and most mammalian cells under rich nutrient conditions (Enenkel, 2014) but relocate to the cytoplasm and form membraneless proteasome storage granules (PSGs) under carbon starvation (Laporte et al., 2008). PSG formation occurs along with reorganization of many other cellular components into granules and foci in response to stress conditions; as with many other cellular granules, PSGs dissipate rapidly once favorable conditions are resumed, with proteasomes reentering the nucleus (Narayanaswamy et al., 2009; Sagot and Laporte, 2019).

Certain types of starvation can also lead to proteasomes being degraded by macroautophagy, likely in a nonselective manner (Li and Hochstrasser, 2020; Marshall et al., 2016; Marshall and Vierstra, 2018; Nemec et al., 2017; Waite et al., 2016). In macroautophagy, portions of the cytoplasm are engulfed by a double-bilayer membrane structure that matures into a closed vesicle called the autophagosome. The autophagosome then fuses with the lysosome (yeast vacuole) membrane, releasing a single large membrane vesicle, the autophagic body (AB), to the lysosome interior. The AB membrane and contents are then broken down by lysosomal hydrolases.

Recently, we have identified an AMPK (AMP-activated protein kinase)-regulated ESCRT (endosomal sorting complex required for transport)-dependent microautophagy pathway of proteasome degradation that occurs concomitantly with PSG formation under low glucose conditions (Li et al., 2019). The ESCRT pathway is a widely deployed mechanism for topologically similar membrane remodeling events such as multivesicular body (MVB) formation, enveloped virus budding, and cell scission (Hurley, 2015). Interestingly, aberrant proteasomes are more likely to be removed by ESCRT-dependent microautophagy than to be stored reversibly in PSGs under these conditions. These findings suggest an important role for AMPK signaling and ESCRT-mediated membrane remodeling in proteasome quality control when glucose levels are low (Li and Hochstrasser, 2020; Segev, 2020).

AMPK and ESCRTs are highly conserved protein complexes participating in diverse cellular processes (Coccetti et al., 2018; Vietri et al., 2020). AMPK is a heterotrimeric complex, composed of one catalytic α subunit and two regulatory subunits, β and γ (Ghillebert, 2011). The kinase is a master regulator of cellular energy homeostasis and is activated by glucose limitation and other stress conditions. In cells, AMPK is regulated by phosphatases and upstream kinases, as well as by autoinhibition of the α subunit (Coccetti et al., 2018). Its activity is altered in many disease states, such as inflammation, diabetes and cancer, and a variety of compounds have been developed to regulate AMPK activity in human disease treatment (Carling, 2017).

The ESCRT factors are part of at least five distinct complexes, ESCRT-0, I, II, III and Vps4 (Schmidt and Teis, 2012). These complexes mediate a series of membrane binding and remodeling steps, starting with target protein recognition and enrichment by ESCRT-0, I and II and leading to membrane remodeling by ESCRT-III and Vps4 (Schmidt and Teis, 2012; Vietri et al., 2020). These steps can occur at different cellular membranes. To understand the roles of ESCRTs in determining the selectivity of proteasome microautophagy, we focused on Vps27, which along with Hse1, forms the Saccharomyces cerevisiae ESCRT-0 complex (Bilodeau et al., 2002). Vps27 is regarded as a “gateway protein” for cargo protein recognition and downstream ESCRT complex assembly (Schmidt and Teis, 2012).

In the current study, we identify a mechanistic role for ESCRT-0 in selective proteasome microautophagy. Specifically, proteasomes in cells subjected to low glucose stress are preferentially modified by low molecular weight (LMW) ubiquitylation of proteasome subunits. Multiple ubiquitylating enzymes contribute to this pathway; we could show that the essential Rsp5 ubiquitin ligase is involved and that the N-lobe noncovalent ubiquitin-binding site in Rsp5 catalytic domain contributes as well. Proteasomes ubiquitylated in this way can be recognized by the ubiquitin-binding domains (UBDs) in the Vps27 subunit of ESCRT-0 and are subsequently cleared through ESCRT-dependent microautophagy. Our data suggest a model in which AMPK functions upstream in the ESCRT-dependent microautophagy pathway under low glucose conditions by regulating Vps27 subcellular trafficking and vacuolar degradation.

## Results

### AMPK promotes Vps27 vacuolar trafficking and degradation in low glucose conditions

To understand how AMPK is involved in ESCRT-dependent microautophagy, we examined steady state levels of Vps27 in various mutants under low glucose (0.025% C) and glucose-free (−C) conditions. Vps27 is known to relocalize from endosomes to the vacuole membrane when cultured yeast cells deplete the glucose supplied in the medium (diauxic shift), and this ESCRT-0 subunit is necessary for microautophagy (Oku et al., 2017). To determine if Vps27 is also degraded in the vacuole during this process, we inactivated vacuolar proteases by deleting the PEP4 and PRB1 genes (Hecht et al., 2014). As shown in Fig. 1A (lane 8 vs. lane 1), disappearance of Vps27 was greatly decreased in this double mutant. Moreover, AMPK was also essential for Vps27 degradation since deletion of either the Snf1 or Snf4 subunits led to strong Vps27 accumulation (Fig. 1A, lanes 2, 3).

**Fig. 1.**
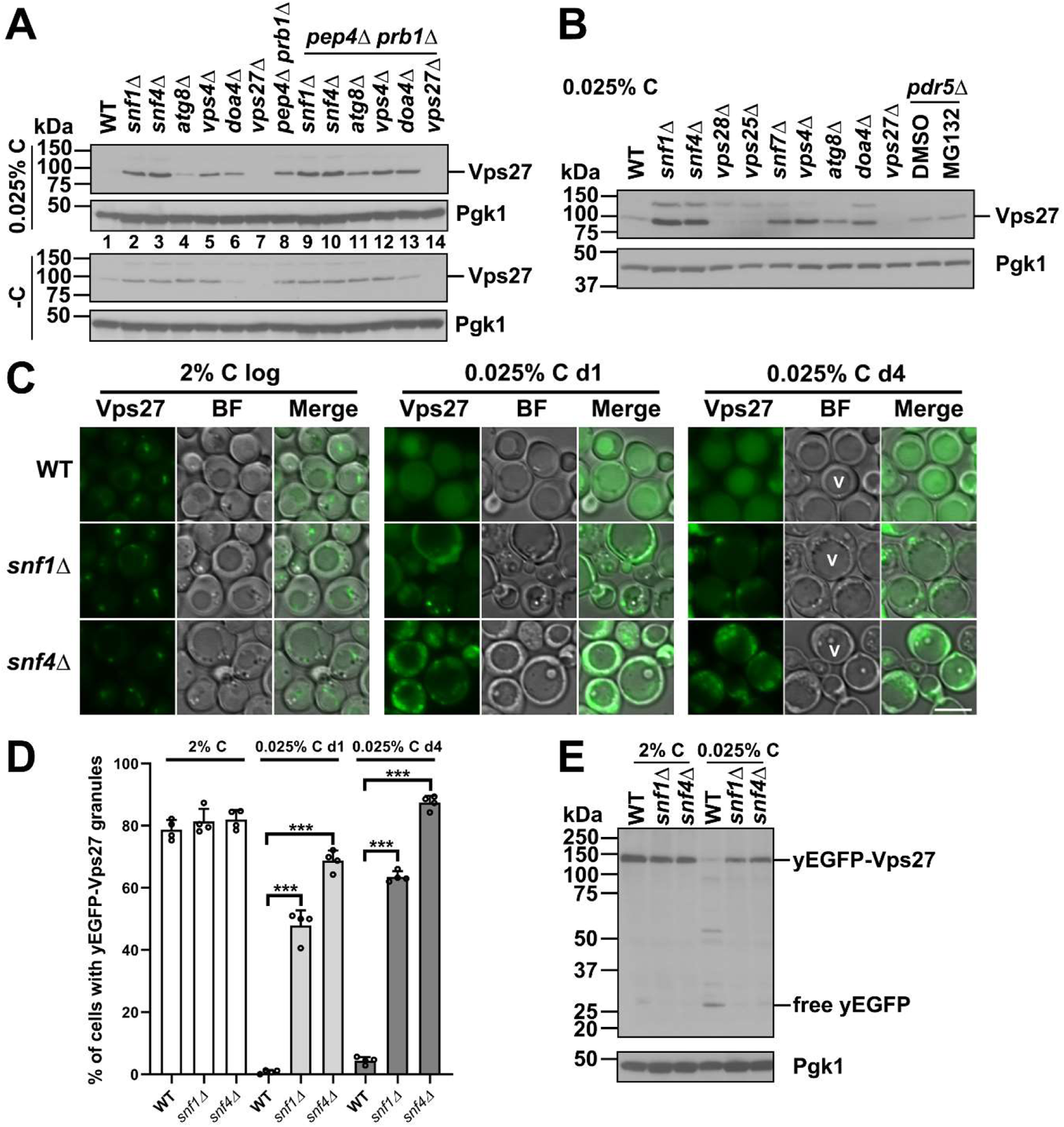
AMPK promotes vacuolar degradation of Vps27 under low glucose conditions. (A) Anti-Vps27 immunoblot analyses in WT, AMPK α subunit mutant snf1Δ and γ subunit mutant snf4Δ, macroautophagy-defective mutant atg8Δ, ESCRT mutants (vps4Δ, vps27Δ), doa4Δ cells, and the corresponding mutant cells in the vacuolar protease-defective mutant pep4Δ prb1Δ background. Cells were harvested from cultures in SC medium containing 0.025% glucose (0.025% C as indicated in the figure panels, or as low glucose medium hereafter) or lacking glucose (-C as indicated in the figure panels) for ~1 d at 30°C. Vps27 was degraded by vacuolar proteases under glucose starvation conditions. (B) Anti-Vps27 immunoblot analyses in WT, snf1Δ, snf4Δ, ESCRT mutants (vps28Δ, vps25Δ, snf7Δ, vps4Δ, vps27Δ), atg8Δ, doa4Δ, or pdr5Δ cells (the last with DMSO or 50 µM MG132 treatments). Cells were harvested from cultures incubated in low glucose medium for ~1 d at 30°C. Vps27 vacuolar degradation was independent of proteasome activity and ESCRT-I (Vps28) or ESCRT-II (Vps25) components. (C) Epifluorescence images of protein expressed from chromosomally integrated yEGFP-VPS27 in WT and AMPK mutant cells. Cells were harvested from cultures at log phase (2% C log) or in low glucose medium for ~1 d or ~4 d at 30°C. Defective trafficking of yEGFP-Vps27 to the vacuole was observed in the mutant cells. BF, bright field. V, vacuole, indicating a representative vacuole in a cell. Scale bar, 5 µm. (D) Percentage of cells with defective trafficking of yEGFP-Vps27 to the vacuole in WT and AMPK mutant cells as analyzed in panel (C). Cells counted at log phase for WT (n=418), snf1Δ (551), and snf4Δ (440); at day 1 low glucose medium for WT (625), snf1Δ (408), snf4Δ (405); and at day 4 low glucose medium for WT (681), snf1Δ (394), snf4Δ (422). Results plotted as mean±sd. ***, P<0.001 (ANOVA single factor analysis comparing AMPK mutant to WT at indicated growth stages). (E) Anti-GFP immunoblot analyses of free yEGFP release from yEGFP-Vps27 in WT and AMPK mutant cells. Cells were from cultures in panel (C) at log phase and low glucose medium for ~1 d at 30°C. yEGFP-Vps27 vacuolar degradation was inhibited in AMPK mutants. Pgk1 served as a loading control. Representative blots and images are shown.

Interestingly, Atg8 and Vps4 made quantitatively distinct contributions to Vps27 degradation in low glucose medium (top), but their loss had similar effects under glucose-free conditions (bottom) (Fig. 1A, lanes 4, 5). Specifically, Vps27 depletion was only weakly impaired in the atg8Δ strain under the former conditions, whereas vps4Δ cells had a greater defect. Atg8 is essential for macroautophagy but is dispensable for microautophagy; under low glucose conditions, defects in microautophagy due to loss of the Vps4 ESCRT factor are much more pronounced (Li et al., 2019). These results suggest that Vps27 degradation occurs primarily via the ESCRT-dependent microautophagy pathway under low glucose conditions.

To investigate the requirement for other ESCRT components in Vps27 degradation, we checked Vps27 steady-state levels in ESCRT-I (Vps28), -II (Vps25), and -III (Snf7) mutants. Vps27 degradation was inhibited in snf7Δ, but not in vps28Δ or vps25Δ cells, suggesting that Vps27 degradation does not go through the canonical MVB pathway, which requires all the major ESCRT factors, including ESCRT-I and ESCRT-II (Fig. 1B). We also tested if degradation of Vps27 depended on proteasome activity under these conditions by treating cells with the MG132 inhibitor. The results in Fig. 1B indicated that proteasomes were not involved in Vps27 depletion.

To visualize whether AMPK mutations affect Vps27 subcellular trafficking, we fused a yeast codon-optimized enhanced green fluorescent protein (yEGFP) to the N-terminus of Vps27 in the context of the normal VPS27 promoter and terminator, and chromosomally integrated the yEGFP-VPS27 fragment into the URA3 locus of vsp27Δ cells. When wild-type (WT) cells were grown in low glucose for one or four days, the yEGFP-Vps27 fusion protein trafficked to the vacuole where it was degraded but left an intact yEGFP cleavage fragment (Fig. 1E); this fragment could also be visualized in the vacuole by yEGFP fluorescence (Fig. 1C). GFP is known to be resistant to vacuolar protease digestion (Klionsky et al., 2021). By contrast, in AMPK mutant cells under the same conditions, yEGFP-Vps27 failed to reach the vacuole and instead concentrated in cytoplasmic granules; these granules underwent what appeared to be rapid docking and separation events (Fig. 1C-1E; highlighted with dashed square in Movie 1, which depicts a snf1Δ cell after one day in low glucose). Therefore, under glucose-limiting conditions, AMPK is required for the trafficking of Vps27 to the vacuole where it would normally be degraded.

Given the punctate cytoplasmic localization of Vps27 in AMPK mutants (Fig. 1C), we examined whether Vps27 cytoplasmic granules colocalized with proteasomal foci under glucose starvation. Interestingly, Vps27 puncta frequently colocalized with proteasomal foci in AMPK mutant cells after four days in low glucose, but not under glucose-free conditions (Fig. S1). Hence, Vps27 might function directly in proteasome quality control by selecting proteasomes for ESCRT-dependent microautophagy in low glucose.

Ubiquitin-binding domains of Vps27 regulate its trafficking and degradation Vps27 contains multiple well-studied functional domains, including the VHS (Vps27, Hrs, and STAM) UBD and two UIMs (ubiquitin-interacting motifs) (Bilodeau et al., 2002) (Fig. 2A). It also has several proline-threonine-alanine-proline-like (PTAPL) motifs for interaction with Vps23 and ESCRT-I complex recruitment (Bilodeau et al., 2003; Katzmann et al., 2003); a FYVE (Fab-1, YOTB, Vps27, and EEA1) domain that binds phosphatidylinositol-3’-monophosphate (PI3P)-rich membranes (Gaullier et al., 1998); and a C-terminal clathrin-binding domain (Bilodeau et al., 2003). We tested whether simultaneous mutation of the Vps27 UBDs, vps27-VHS-UIM (Ren and Hurley, 2010), affected vacuolar degradation of Vps27 in low glucose. Notably, few apparent yEGFP cleavage products were detected, yet yEGFP-vps27-VHS-UIM loss was still apparent (Fig. 2B). This suggested that vps27-VHS-UIM degradation might involve a pathway other than autophagy. Indeed, disappearance of the mutant protein, unlike the WT version, was inhibited in MG132-treated pdr5Δ cells, and this occurred regardless of whether Vps27 was fused to yEGFP (Fig. 2B, E).

**Fig. 2.**
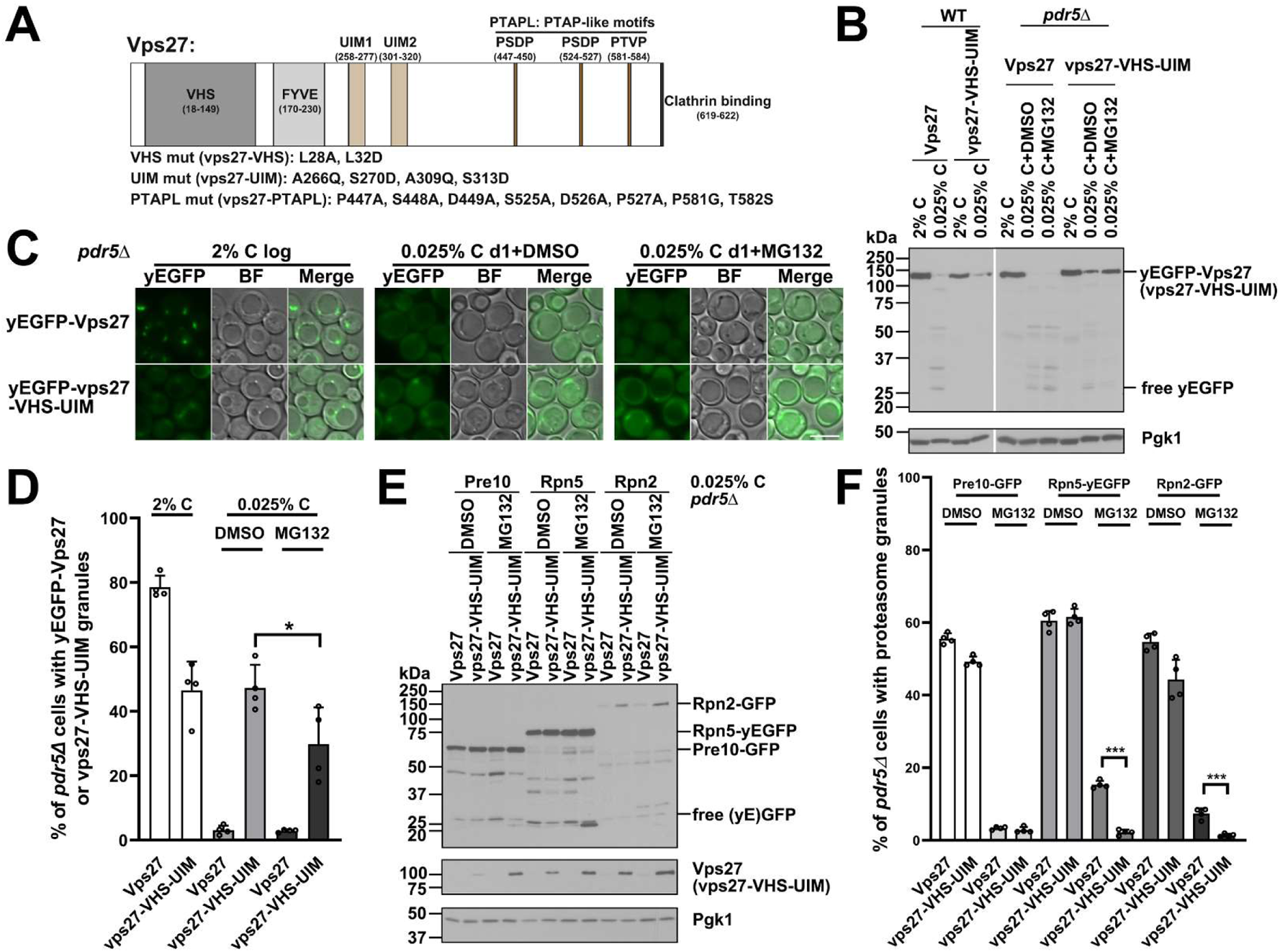
Mutation of Vps27 UBDs impairs Vps27 and proteasome trafficking and degradation. (A) Schematic of Vps27 functional domains and the indicated vps27 mutations used in this study. (B) Anti-GFP immunoblot analyses of free yEGFP release from yEGFP-Vps27 and an ubiquitin binding-defective Vps27 mutant (yEGFP-vps27-VHS-UIM) in WT and pdr5Δ cells. Endogenous VPS27 was deleted in all the strains. Cells were harvested from cultures in exponential phase or in low glucose medium for ~1 d at 30°C. Additionally, pdr5Δ cells were treated with DMSO or 50 µM MG132. yEGFP-vps27-VHS-UIM degradation was affected by proteasome activity in cells under low glucose conditions. (C) Fluorescence images of yEGFP-Vps27 and yEGFP-vps27-VHS-UIM in pdr5Δ cells from panel (B). Ubiquitin-binding domains (UBDs) of Vps27 contributed to Vps27 trafficking and its degradation by autophagy and proteasomes. BF, bright field. Scale bar, 5 µm. (D) Percentage of pdr5Δ cells with yEGFP-Vps27 and yEGFP-vps27-VHS-UIM foci. Cells were from cultures in panel (B). Cells counted at log phase: VPS27 (n=483), vps27-VHS-UIM (540; at day 1 low glucose medium with DMSO: VPS27 (587), vps27-VHS-UIM (599); and at day 1 low glucose medium with MG132: VPS27 (481), vps27-VHS-UIM (600). Results plotted as mean±sd. *, P<0.05 (ANOVA single factor analysis comparing MG132- to DMSO only-treated vps27-VHS-UIM cells). (E) Anti-GFP immunoblot analyses of free (yE)GFP release and proteasome fragmentation (proteolysis products in between full length GFP fusion protein and free GFP) of Pre10-GFP (a CP subunit), Rpn5-yEGFP (a lid subunit), and Rpn2-GFP (a base subunit) in pdr5Δ cells with VPS27 or vps27-VHS-UIM alleles controlled by the natural VPS27 promoter and terminator and integrated at the URA3 locus. Endogenous VPS27 gene was deleted in all the strains. Cells were harvested from cultures grown in low glucose medium containing DMSO or 50 µM MG132 for ~1 d at 30°C. Proteasome subunit fragmentation was enhanced by MG132 treatment but inhibited by vps27-VHS-UIM. Middle panel: Vacuolar degradation of untagged vps27-VHS-UIM was inhibited in cells treated with MG132, consistent with the effect seen with yEGFP fused vps27-VHS-UIM. (F) Percentage of pdr5Δ cells with proteasomal puncta from cultures in panel (E). Cells counted from cultures treated with DMSO and expressing Pre10-GFP (402, 487 for VPS27 and *vps27-VHS-UIM*, respectively); Rpn5-yEGFP: (632, 517) or Rpn2-GFP (614, 412), and from cultures with treated with MG132 and expressing Pre10-GFP (444, 602); Rpn5-yEGFP (434, 327); or Rpn2-GFP (526, 785). PSG formation was inhibited by MG132 treatment, and RP-containing PSG formation was further suppressed in MG132-treated *vps27-VHS-UIM* cells. Results plotted as mean±sd. ***, P<0.001 (ANOVA single factor analysis comparing *vps27-VHS-UIM* to VPS27 from cultures with MG132 treatment). Representative blots and images are shown.

Membrane association and vacuolar trafficking of yEGFP-vps27-VHS-UIM also appeared to be disrupted (Fig. 2C, D). In cells grown in rich medium, WT yEGFP-Vps27 concentrated in small foci or granules, which are assumed to be endosomes (Hatakeyama et al., 2019; Katzmann et al., 2003), whereas the mutant derivative displayed a more diffuse pattern, which correlates with cargo protein-sorting defects (Ren and Hurley, 2010). In low glucose, WT yEGFP-Vps27 did not accumulate appreciably within cytoplasmic granules even in the presence of MG132; proteasome inactivation slightly reduced vps27-VHS-UIM granule staining (Fig. 2C, D). These results confirmed that the Vps27 UBDs, and presumably ubiquitin binding, are needed for efficient Vps27 recruitment to endosomes and thus proper cargo sorting, as previously reported (Ren and Hurley, 2010). They also suggest that these domains modulate the interaction of Vps27 with proteasomes during carbon limitation.

Vps27, as well as components of all the other ESCRT complexes, are essential for proteasome microautophagy during low-glucose stress (Li et al., 2019). This can be followed by the fragmentation of proteasome subunits into GFP-sized and larger fragments; these more complex fragmentation patterns are specifically seen when microautophagy is active, although the basis of this is unclear (Li et al., 2019). They nevertheless provide a useful signature for microautophagy-mediated proteasome degradation and likely other substrates such as Vps27 itself, e.g., Fig. 2B.

It is striking that despite the sharp drop in overall Vps27 levels during carbon limitation, Vps27-dependent proteasome microautophagy, as measured by subunit fragmentation, continued (Fig. 2E). In the vps27-VHS-UIM mutant, proteasome fragmentation was modestly reduced under these conditions, while as noted above, the vps27-VHS-UIM protein itself was stabilized. Inhibition of proteasome activity generally enhances detection of fragmented proteasome subunits (Li et al., 2019), so we determined if MG132 still affected the degree of subunit fragmentation in the vps27-VHS-UIM mutant. As shown in Fig. 2E, in glucose-starved proteasome-inhibited cells, a moderate reduction in proteasome subunit fragmentation was seen compared to similarly inhibited WT cells. At the same time, PSG formation, in particular that of RP-containing PSGs, was significantly reduced in MG132-treated vps27-VHS-UIM cells accompanied by enhanced nuclear proteasomes (Figs. 2F, S2). From these results, we hypothesize that proteasomes, particularly aberrant or inactive ones, are recognized by Vps27 for microautophagy during glucose starvation and that this requires the Vps27 UBDs for full efficacy.

### Proteasome subunit fragmentation is ubiquitin dependent

To begin testing this hypothesis, we checked whether ubiquitin affected proteasome subunit degradation, since ubiquitylation plays an important role in cargo protein recognition by ESCRTs (Shields and Piper, 2011). Doa4 is a DUB enzyme that participates in the ESCRT pathway by removing ubiquitin from ubiquitylated cargos (Amerik et al., 2000), while deleting DOA4 disrupts ubiquitin homeostasis and causes ubiquitin depletion from cells (Swaminathan et al., 1999). The microautophagy-specific proteasome subunit fragmentation pattern was lost in the doa4Δ mutant, while generation of free GFP from GFP-tagged subunits increased (Fig. 3A), presumably due to concomitant macroautophagy (Li et al., 2019). The WT fragmentation pattern was restored by ectopically expressing ubiquitin (Fig. 3A), suggesting microautophagy of proteasomes is ubiquitin dependent. As expected, Doa4 DUB activity is required for subunit fragmentation as the Doa4 catalytically dead mutant doa4-C571S (Papa and Hochstrasser, 1993) failed to rescue the defect (Fig. 3B).

**Fig. 3.**
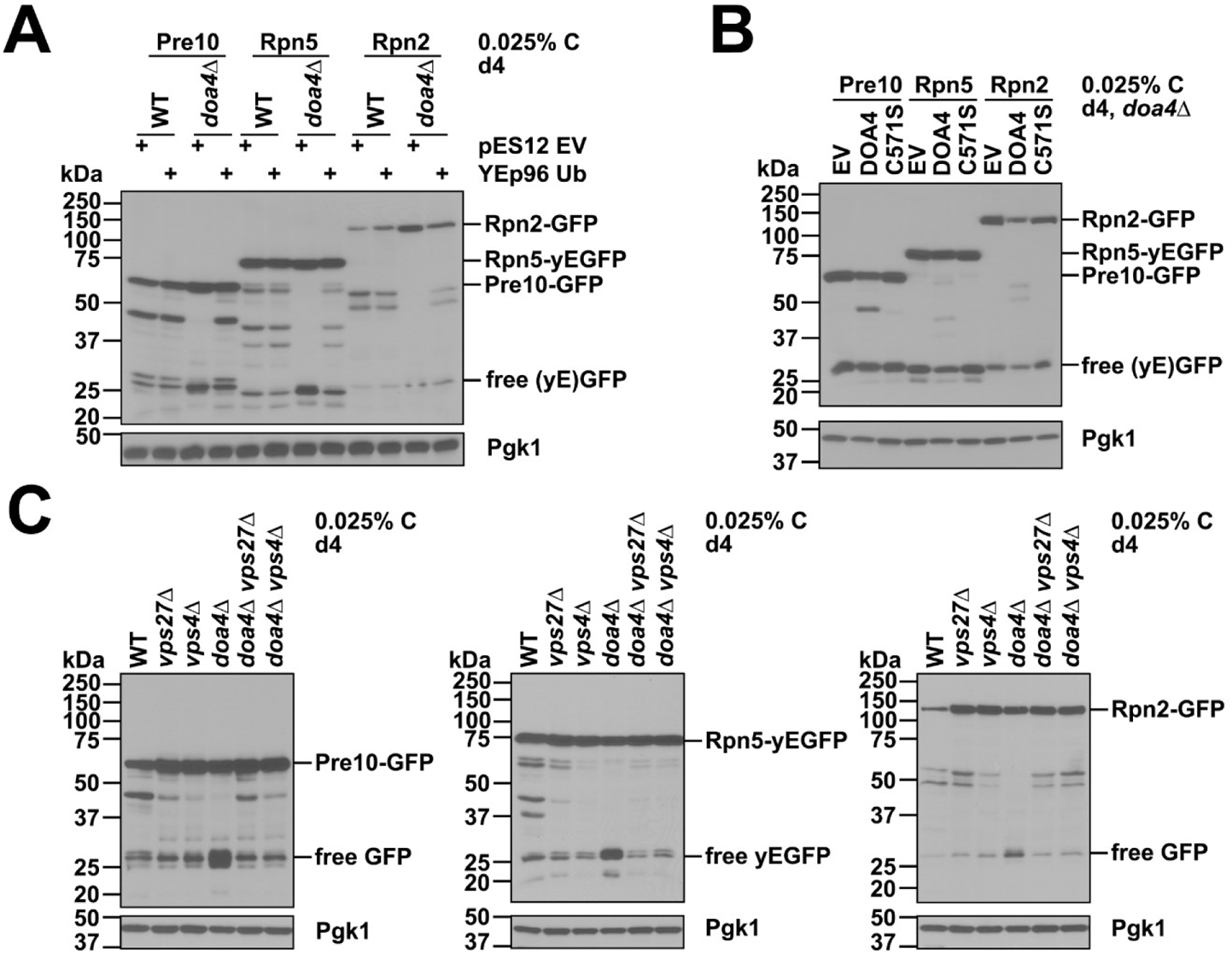
Microautophagy-dependent proteasome subunit fragmentation is ubiquitin dependent. (A) Anti-GFP immunoblot analyses of free (yE)GFP release and proteasome fragmentation of Pre10-GFP, Rpn5-yEGFP, and Rpn2-GFP in WT and doa4Δ cells carrying either pES12 (empty vector) or YEp96 for ubiquitin expression under the copper-inducible CUP1 promoter (basal expression, no copper added). Overexpressing ubiquitin suppressed the loss of proteasome fragmentation in doa4Δ cells. (B) Anti-GFP immunoblot analyses of free (yE)GFP release and proteasome fragmentation of Pre10-GFP, Rpn5-yEGFP, and Rpn2-GFP in doa4Δ cells expressing pRS316-based alleles: empty vector (EV), wild type DOA4, or catalytically inactive mutant doa4-C571S under the control of the DOA4 native promoter. Doa4 DUB activity was required for microautophagy-dependent proteasome fragmentation. (C) Anti-GFP immunoblot analyses of free (yE)GFP release and proteasome fragmentation of Pre10-GFP, Rpn5-yEGFP, and Rpn2-GFP in the indicated strains. ESCRT gene deletions in doa4Δ that suppress ubiquitin depletion could also rescue proteasome fragmentation in doa4Δ cells. Representative blots are shown.

Mutations of ESCRT components interact genetically with doa4Δ and suppress the ubiquitin depletion phenotype in cells lacking active Doa4 (Swaminathan et al., 1999). The microautophagy-specific proteasome fragmentation pattern was restored in doa4Δ vps27Δ and doa4Δ vps4Δ cells to the levels of the vps27Δ and vps4Δ single deletions (Fig. 3C); this partial suppression was expected because these ESCRT components are themselves necessary for normal proteasome fragmentation in low glucose conditions (Li et al., 2019). AMPK is also necessary for proteasome microautophagy (Li et al., 2019), but loss of AMPK, unlike loss of the ESCRT factors, did not suppress the fragmentation defects of doa4Δ cells, although it did suppress the hyperaccumulation of free GFP (Fig. S3). This could be due to the more penetrant effect on proteasome fragmentation of AMPK (Li et al., 2019). Overall, these results suggest that ubiquitin is important for ESCRT-mediated proteasome sorting and degradation by microautophagy.

### LMW ubiquitylation of proteasomes is observed in ESCRT mutants

Using a nanobody to GFP, we directly examined proteasome ubiquitylation by co-immunoprecipitation (co-IP) from extracts of cells expressing GFP-tagged proteasomes. Interestingly, we observed a distinct low molecular weight (LMW) ubiquitylation pattern of proteasome subunits in ESCRT mutants along with a reduction in high molecular weight (HMW) conjugates; the latter are likely to be noncovalently bound ubiquitylated proteasome substrates (Figs. 4, S4A). A broad range of LMW ubiquitylated species also accumulated in total cell lysates of ESCRT mutants; however, we attributed this to the known accumulation of Lys63-linked ubiquitin chains conjugated to cargo proteins that is caused by the block in the MVB pathway in these ESCRT mutants (Lauwers et al., 2009). Importantly, no ubiquitin-K63 chain conjugates were detected in the proteasome co-IPs (Fig. S4B); consistent with this, in the IP samples, we detected a small number of ubiquitin-ubiquitin linkages by mass spectrometry (MS) and all involved K48. Proteasome ubiquitylation was also found to be higher following large-scale affinity purifications from ESCRT mutants expressing a 3xFLAG-tagged Rpn11 RP subunit (Fig. S5). Proteasomes that were salt washed (Marshall et al., 2016) retained the LMW ubiquitylated species (Fig. S4C). Finally, besides ubiquitin, proteasome subunits and proteasome-associated proteins such as RP assembly chaperones were the dominant proteins identified by MS analysis from the immunoprecipitated proteasomes from WT and ESCRT mutants (Fig. S6, Table S1), also arguing against ubiquitylated substrates bound to proteasomes being major contributors to the LMW ubiquitylation species in ESCRT mutants. Overall, these results support the idea that the LMW ubiquitylated species are primarily ubiquitin-modified proteasome subunits that accumulate in ESCRT mutants under low glucose conditions.

**Fig. 4.**
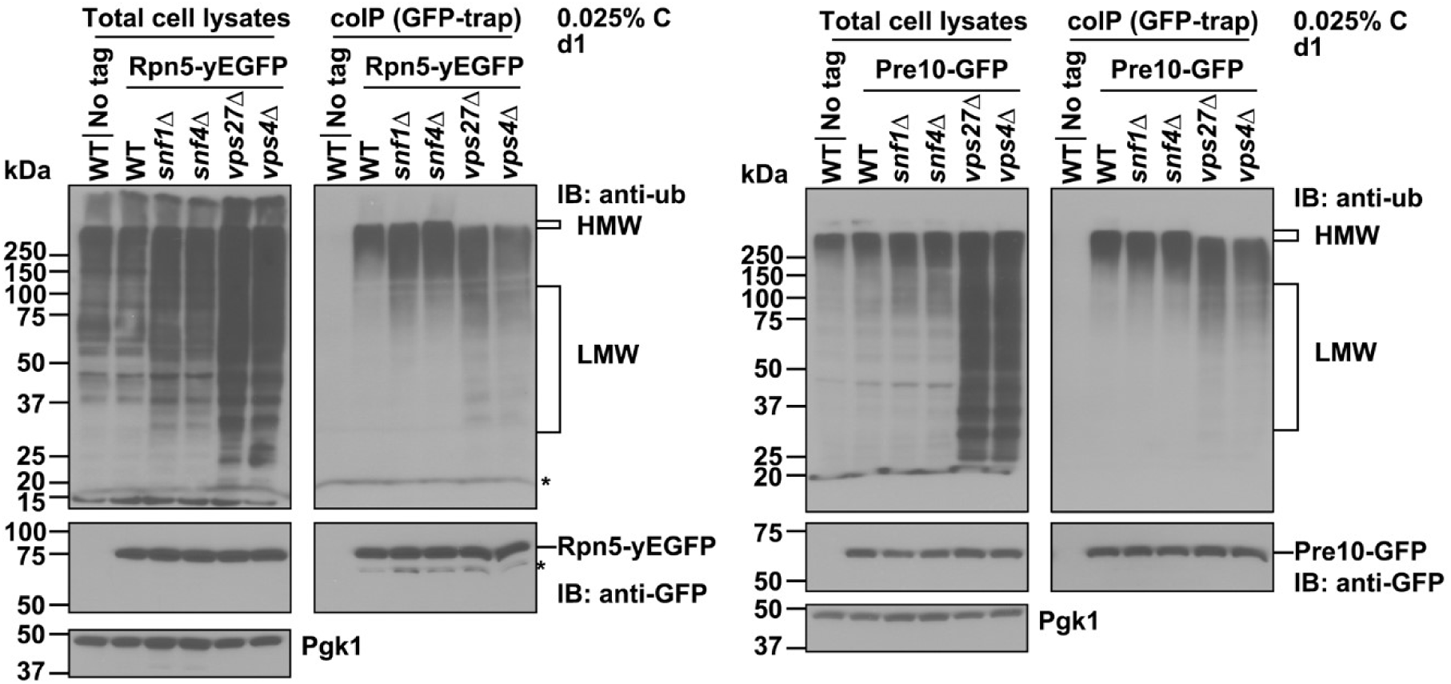
Low molecular weight (LMW) ubiquitylated proteasome species accumulate in ESCRT mutants. Co-immunoprecipitation (co-IP) analyses of ubiquitylated proteasome species in WT cells or the indicated expressing Rpn5-yEGFP (left panel) or Pre10-GFP (right panel). LMW ubiquitylated proteasome species accumulated in ESCRT mutants under low glucose conditions. GFP-trap agarose resin was used to immunoprecipitate proteasomes. Cells were harvested from cultures grown in low glucose medium for ~1 d at 30°C. *, nonspecific bands. HMW, high molecular weight. Representative blots are shown.

### Aberrant proteasomes are subject to LMW ubiquitylation and cleared through ESCRT-0

To test the hypothesis that aberrant or incompletely assembled proteasomes are preferred substrates for ubiquitylation, we performed proteasome ubiquitylation analyses in proteasome mutants (pre9Δ and sem1Δ) that bear high amounts of aberrant proteasomes (Tomko Jr and Hochstrasser, 2014; Velichutina et al., 2004) as well as in MG132-treated pdr5Δ cells containing catalytically inhibited proteasomes. As predicted, LMW ubiquitylated proteasome species were enriched in pre9Δ and MG132-treated pdr5Δ cells (Fig. 5A). This was not the case for sem1Δ cells; this mutation limits 26S proteasome egress from the nucleus and prevents PSG accumulation in low glucose (Li et al., 2019), suggesting proteasome subunit ubiquitylation occurs after nuclear export. Conversely, ubiquitylation of proteasomes was almost abolished in doa4Δ cells, which have very little free ubiquitin (Fig. 5A), in line with the role of ubiquitin in proteasome fragmentation (Fig. 3). These results are consistent with aberrant proteasomes being more strongly modified by LMW ubiquitylation.

**Fig. 5.**
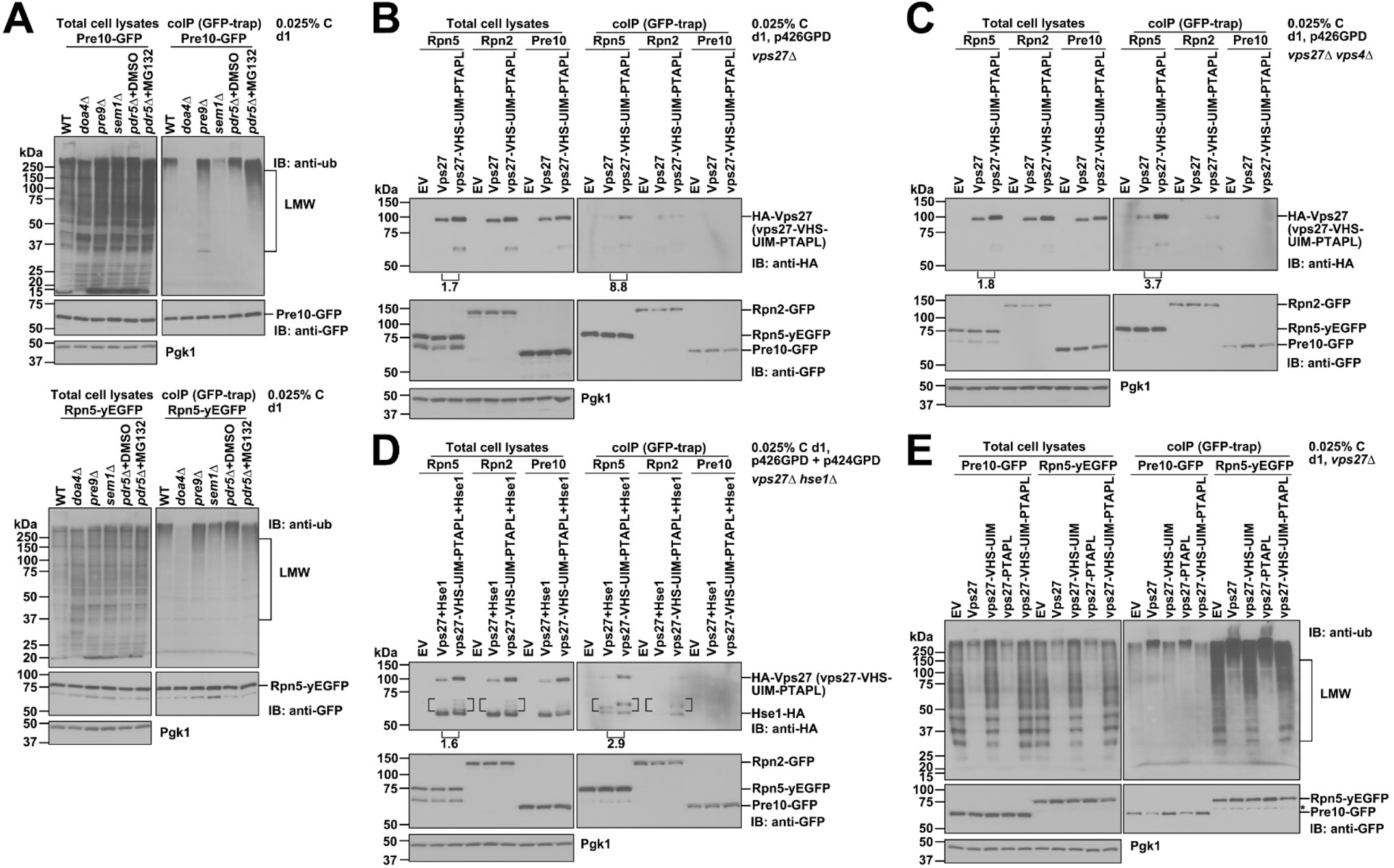
LMW ubiquitylation of aberrant proteasomes and their removal by ESCRT-0. (A) Co-IP analyses of ubiquitylated proteasome species in WT, doa4Δ, proteasome mutants (pre9Δ, sem1Δ), and MG132-treated cells expressing Pre10-GFP (top panel) and Rpn5-yEGFP (bottom panel). LMW ubiquitylated proteasome species accumulated in pre9Δ and MG132-treated cells. (B, C) Co-IP analyses of HA-Vps27 and HA-vps27-VHS-UIM-PTAPL with proteasomes in vps27Δ (B) and vps27Δ vps4Δ (C) cells expressing Rpn5-yEGFP, Rpn2-GFP, or Pre10-GFP. Relative abundance of full-length HA-vps27-VHS-UIM-PTAPL and HA-Vps27 was calculated by normalizing to full-length Rpn5-yEGFP in total cell lysates or co-IP; the ratio of HA-vps27 mutant to HA-Vps27 is noted in panels. Vps27 interacted with proteasome RP but not CP under low glucose conditions, and this interaction increased when VPS4 was deleted; vps27-VHS-UIM-PTAPL was stabilized due to stalled microautophagy. (D) co-IP analyses of HA-tagged ESCRT-0 subunits (Vps27, Hse1) and proteasomes as in panel (C). The protein abundance of ESCRT-0 including full length and intermediates or breakdown products (as indicated with brackets in the figure panels) in between HA-Vps27 or HA-vps27-VHS-UIM-PTAPL and Hse1-HA was quantified as in panel (C). (E) co-IP analyses of ubiquitylated proteasome species in vps27Δ cells bearing chromosomally integrated pRS306 (EV) or the indicated VPS27 alleles. LMW ubiquitylated proteasome species accumulated in ubiquitin binding-defective Vps27 mutants. Cells were harvested from cultures grown in low glucose medium for ~1 d at 30°C. Protein band intensity was quantified with ImageJ. Representative blots are shown.

Next, we tested whether ESCRT-0 subunit Vps27 interacted with proteasomes. Given that endogenous Vps27 was also significantly degraded by microautophagy in low glucose (Fig. 1) and that ESCRT-0-proteasome interaction is likely transient, we made constructs for constitutive overexpression of HA-tagged Vps27 and a mutant vsp27-VHS-UIM-PTAPL construct expressed from the strong GPD promoter to facilitate detection of Vps27-proteasome interaction. The vps27-VHS-UIM-PTAPL mutant protein (Fig. 2A) was designed to reduce or eliminate the ubiquitin-binding ability by mutating the Vps27 UBDs and simultaneously disturbing its association with downstream ubiquitin-binding ESCRT complexes (Bilodeau et al., 2002; Bilodeau et al., 2003; Katzmann et al., 2003). Multiple UBDs have been identified in ESCRT components, including Hse1 and Vps27 in ESCRT-0, Vps23 and Mvb12 in ESCRT-I, and Vps36 in ESCRT-II (Schmidt and Teis, 2012). The Vps27 PTAPL motifs interact with Vps23 for ESCRT-I recruitment, thereby increasing sorting efficiency of ubiquitylated cargo proteins (Bilodeau et al., 2003; Katzmann et al., 2003). With GPD-driven expression, HA-Vps27 and HA-vps27-VHS-UIM-PTAPL could be detected in cell lysates from vps27Δ cells grown under low glucose conditions (Fig. 5B). No ATP was added to the IP lysis buffer, which favored CP-RP dissociation so that any preference of Vps27 for binding CP or RP could be determined.

Vps27 interacted with the proteasome RP complex, as demonstrated by its co-IP by anti-GFP antibodies when RP subunits Rpn5 or Rpn2 were tagged with GFP; binding was not detected with the CP, using CP subunit Pre10-GFP for co-IP (Fig. 5B). Surprisingly, the vps27-VHS-UIM-PTAPL mutant co-precipitated more RP complex than did WT Vps27, but there was also more of the mutant ESCRT subunit in the total lysates (Fig. 5B). This suggests that Vps27 can likely interact with the RP in ways that are independent of RP ubiquitylation and also that vps27-VHS-UIM-PTAPL may be less susceptible to degradation, possibly by stalling microautophagy. Consistent with this interpretation, the difference between WT and mutant Vps27 binding to the RP complex was similar in vps4Δ cells (Fig. 5C), which cannot carry out microautophagy (Li et al., 2019).

Vps27 interacts with Hse1 to form the ESCRT-0 complex (Bilodeau et al., 2003; Ren and Hurley, 2010). Hse1 is unable to bind ubiquitin in vitro in the absence of Vps27 (Bilodeau et al., 2002). We checked whether overproduced Hse1 was sufficient to interact with proteasomes in vivo. Using GPD-driven Hse1-HA expression in hse1Δ cells, we did not detect any interactions with proteasomes, nor did levels of LMW ubiquitylated proteasome species change (Fig. S7). However, when both Vps27 and Hse1 were overproduced, Hse1 co-precipitated with the RP along with Vps27 (Fig. 5D), suggesting that the ESCRT-0 complex binds the RP via its Vps27 subunit.

We predicted that a Vps27 mutant, such as vps27-VHS-UIM-PTAPL that disturbed the ubiquitin-binding ability of ESCRT components, should accumulate (aberrant) proteasomes that are modified by LMW ubiquitylation under glucose-limited conditions. To reconstitute endogenous levels of Vps27, we created a set of strains with either WT VPS27 or different vps27 mutant alleles (vps27-VHS-UIM, -PTAPL, -VHS-UIM-PTAPL) integrated at the chromosomal URA3 locus; the alleles were under the control of the natural VPS27 promoter and terminator in vps27Δ cells that also expressed GFP-tagged proteasome subunits (Pre10 or Rpn5). Integration of the empty vector served as a negative control. Based on proteasome ubiquitylation analysis, LMW ubiquitylated proteasome species accumulated in the vps27-VHS-UIM-PTAPL and vps27-VHS-UIM strains but not in VPS27 or vps27-PTAPL cells (Fig. 5E). This suggests that proteasomes modified by LMW ubiquitylation accumulate when these ubiquitylated forms cannot be recognized by ESCRT-0.

It should be noted that these mutants will also block other ESCRT-0-dependent mechanisms such as the endosomal-MVB pathway (Bilodeau et al., 2003; Katzmann et al., 2003; Ren and Hurley, 2010). A distinctive ladder of ubiquitylated species accumulated in the mutants, as seen in the total lysates in Fig. 5E. These may be K63-linked ubiquitin oligomers that are important in the MVB pathway but appear to play little or no role in ubiquitylation of proteasomes (Fig. S4B). Because of their abundance, a fraction may nevertheless bind to proteasomes under the conditions used for the experiment in Fig. 5E.

### Ubiquitin-ESCRT binding is essential for ILV formation and proteasome microautophagy

We next examined whether the above Vps27 mutants also affected proteasome trafficking and degradation in the vacuole. We used the same set of yeast strains (Fig. 5E) and checked proteasome fragmentation (as a biochemical signature of microautophagy) and trafficking. Agreeing well with the data from vps27-VHS-UIM pdr5Δ cells (Fig. 2E), a minor proteasome fragmentation defect was observed in vps27-VHS-UIM cells (Fig. 6A). As predicted from proteasome ubiquitylation (Fig. 5E), vps27-PTAPL cells did not have any defects in proteasome fragmentation; by contrast, a much stronger defect in proteasome fragmentation was detected in vps27-VHS-UIM-PTAPL cells, particular after four days of glucose deprivation (Fig. 6A), indicating the vps27-VHS-UIM-PTAPL mutations disrupted ESCRT-dependent microautophagy of proteasomes to the same extent as vps27Δ cells.

**Fig. 6.**
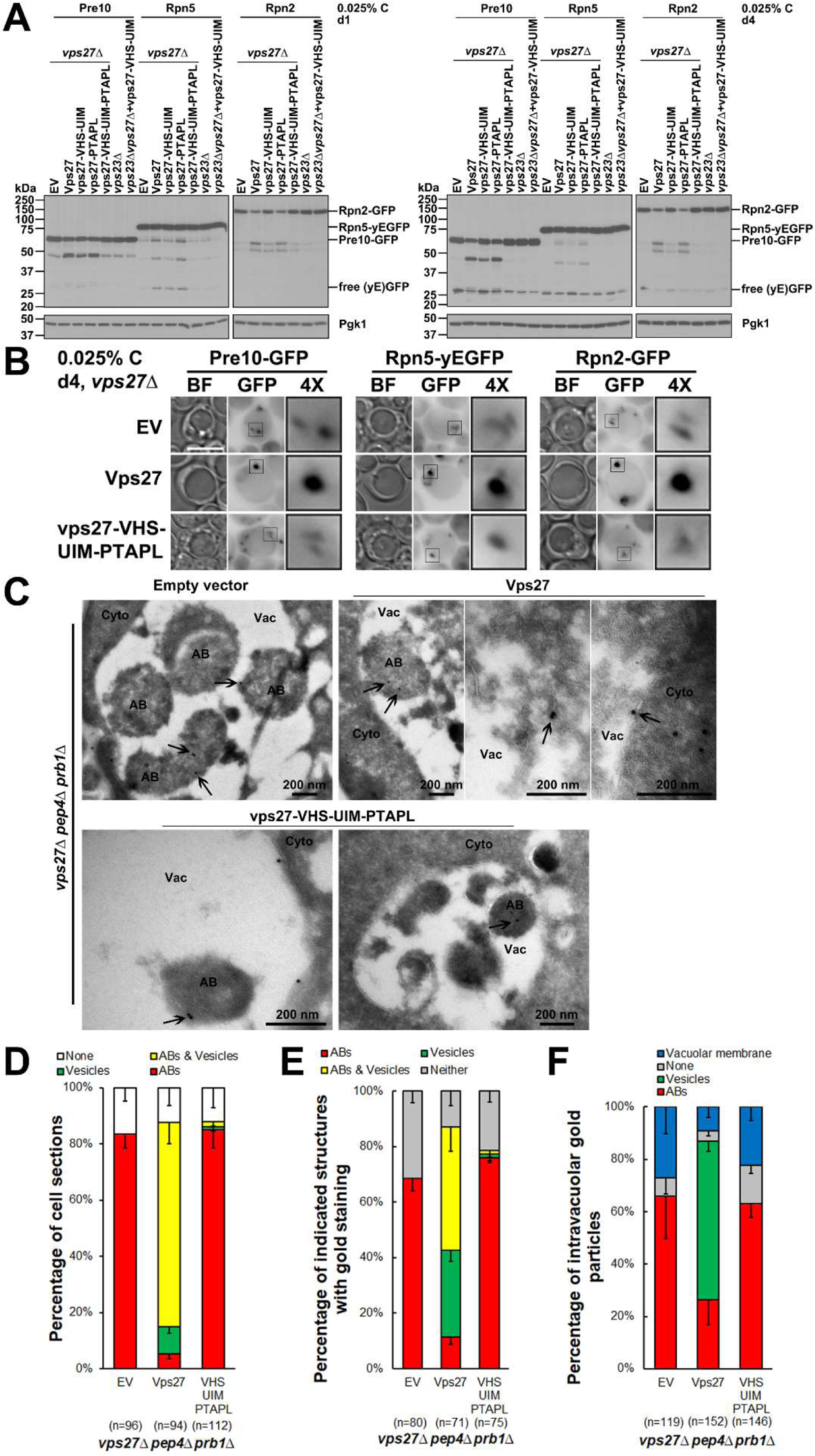
Ubiquitin binding by ESCRT components is essential for proteasome microautophagy and intralumenal vesicle (ILV) formation. (A) Anti-GFP immunoblot analyses of Pre10-GFP, Rpn5-yEGFP, and Rpn2-GFP in vps27Δ cells harboring chromosomally integrated empty vector (EV, pRS306) and the indicated VPS27 alleles in the strains shown. Cells were harvested from cultures grown in low glucose medium for ~1 d and 4 d at 30°C. Representative blots are shown. (B) Epifluorescence images of Pre10-GFP, Rpn5-yEGFP, and Rpn2-GFP in vps27Δ cells bearing the indicated chromosomally integrated VPS27 alleles. Cells were from cultures in panel (A) after ~4 d in low glucose. Proteasome puncta were observed in EV and vps27-VHS-UIM-PTAPL cells, indicating accumulation of aberrant proteasomes. 4×, 4-fold enlargement of the boxed regions. Scale bar, 5 µm. Representative images are shown. (C) Cryo-immunogold electron micrographs of proteasomes in vps27Δ pep4Δ prb1Δ cells expressing chromosomally integrated EV, VPS27, or vps27-VHS-UIM-PTAPL. Cells were grown in low glucose for ~4 d at 30°C. Sections were immunolabeled with anti-CP primary antibody and gold-conjugated Protein A (black arrows). Cyto, cytoplasm; Vac, vacuole; AB, autophagic body. (D) Percentage of cell sections with the indicated intravacuolar structures (ABs, vesicles, ABs & vesicles, none) in the strains used for panel (C). (E) Percentage of the indicated structures with gold particle staining in the cells used for panel (C). (E) Percentage of intravacuolar gold particles staining either ABs or vesicles or the limiting vacuolar membrane in the cells used for panel (C). “n” represents cell sections counted in panels (D) and (E), and intravacuolar gold particles counted in panel (F).

As noted above, Vps23 (ESCRT-I) directly associates with Vps27 by binding its PTAPL motifs (Bilodeau et al., 2003; Katzmann et al., 2003). We created a vps27-VHS-UIM vps23Δ strain, which retains the PTAPL elements in Vps27, and determined whether this mutant mimicked the proteasome microautophagy defect of vps27-VHS-UIM-PTAPL VPS23 cells. Interestingly, an additive defect in proteasome fragmentation was observed in vps27-VHS-UIM vps23Δ as compared to the individual vps23Δ and vps27-VHS-UIM-PTAPL mutants, especially in low glucose conditions, at early times (Fig. 6A, left panel, e.g. lane 7 vs. lanes 5, 6 in the set of Rpn2-GFP strains), suggesting that these ESCRT components function collaboratively to recognize proteasomes for microautophagy.

Cytological analyses by fluorescence and immunogold electron microscopy further demonstrated equivalent defects in proteasome trafficking and degradation in vps27-VHS-UIM-PTAPL and vps27Δ cells. Rather than forming spherical and bright PSGs as in VPS27 (WT) cells, irregular and faint proteasome puncta were observed in the cytoplasm, particularly at the perivacuolar region of vps27-VHS-UIM-PTAPL and EV (vps27Δ) cells (Fig. 6B), suggesting an accumulation of aberrant proteasomes in the glucose-limited cells. To preserve autophagic body (AB) and intralumenal vesicle (ILV) structures generated by macroautophagy and microautophagy, respectively, under low glucose stress, the chromosomally integrated VPS27 and vps27-VHS-UIM-PTAPL alleles (at the URA3 locus) were placed in a vps27Δ pep4Δ prb1Δ background. Although ILV formation in the vacuole was previously observed with a Vps27-Hse1 complex lacking any intact UIMs (Bilodeau et al., 2002), ILV formation was blocked and only ABs were observed in vps27-VHS-UIM-PTAPL and vps27Δ (EV) cells, while both ILVs and ABs were observed in VPS27 cells (Fig. 6C, D). This phenotype is reminiscent of the microautophagy defect in vps4Δ cells (Li et al., 2019). Proteasome distribution in the vacuole was followed by anti-CP antibodies labeled with gold beads (black arrows in Fig. 6C). A comparable proteasome distribution pattern was found in vps27-VHS-UIM-PTAPL and vps27Δ (EV) cells (Fig. 6E, F). The results suggest that the ubiquitin-binding ability of ESCRTs is essential for ILV formation and thus proteasome microautophagy.

Rsp5 ubiquitin ligase is involved in proteasome ubiquitylation

Given the intriguing ubiquitylation pattern of proteasomes in ESCRT mutants, we sought to find the ubiquitin ligase(s) responsible for their ubiquitylation. Rsp5 is the principal ligase involved in cargo protein ubiquitylation in the ESCRT-dependent endosomal/MVB pathway of protein targeting to the vacuole and is also known to interact with ESCRT-0 (MacDonald et al., 2020). Rsp5 belongs to the NEDD4 family of HECT (homologous to E6AP C-terminus) ligases (Shah and Kumar, 2021). NEDD4 ligases share an N-terminal lipid-interacting C2 domain, 1-4 central WW protein interaction motifs (Chen and Sudol, 1995), and a large C-terminal catalytic HECT domain; the HECT domain bears an N-lobe with a noncovalent ubiquitin-binding site and a C-lobe with a covalent (catalytic) ubiquitin-binding site (Finley et al., 2012; Shah and Kumar, 2021).

We tested whether Rsp5 contributed to proteasome ubiquitylation under low glucose conditions. Rsp5 is essential for cell viability in yeast, so we performed the analysis in a temperature sensitive (ts) mutant, rsp5-1, which has an L733S mutation that causes dysfunction of Rsp5 at non-permissive temperatures (Wang et al., 1999). Proteasome ubiquitylation was inhibited in rsp5-1 at elevated temperature (30°C) but was comparable to WT at room temperature (RT, ~24°C) (Fig. 7A). The defects in proteasome ubiquitylation correlated with reduced proteasome subunit fragmentation in rsp5-1 (Fig. S8). Fragmentation of RP subunits Rpn5 and Rpn2 was more sensitive to defective Rsp5 compared to the Pre10 CP subunit, suggesting the RP is the main ubiquitylation target for initiating proteasome microautophagy.

**Fig. 7.**
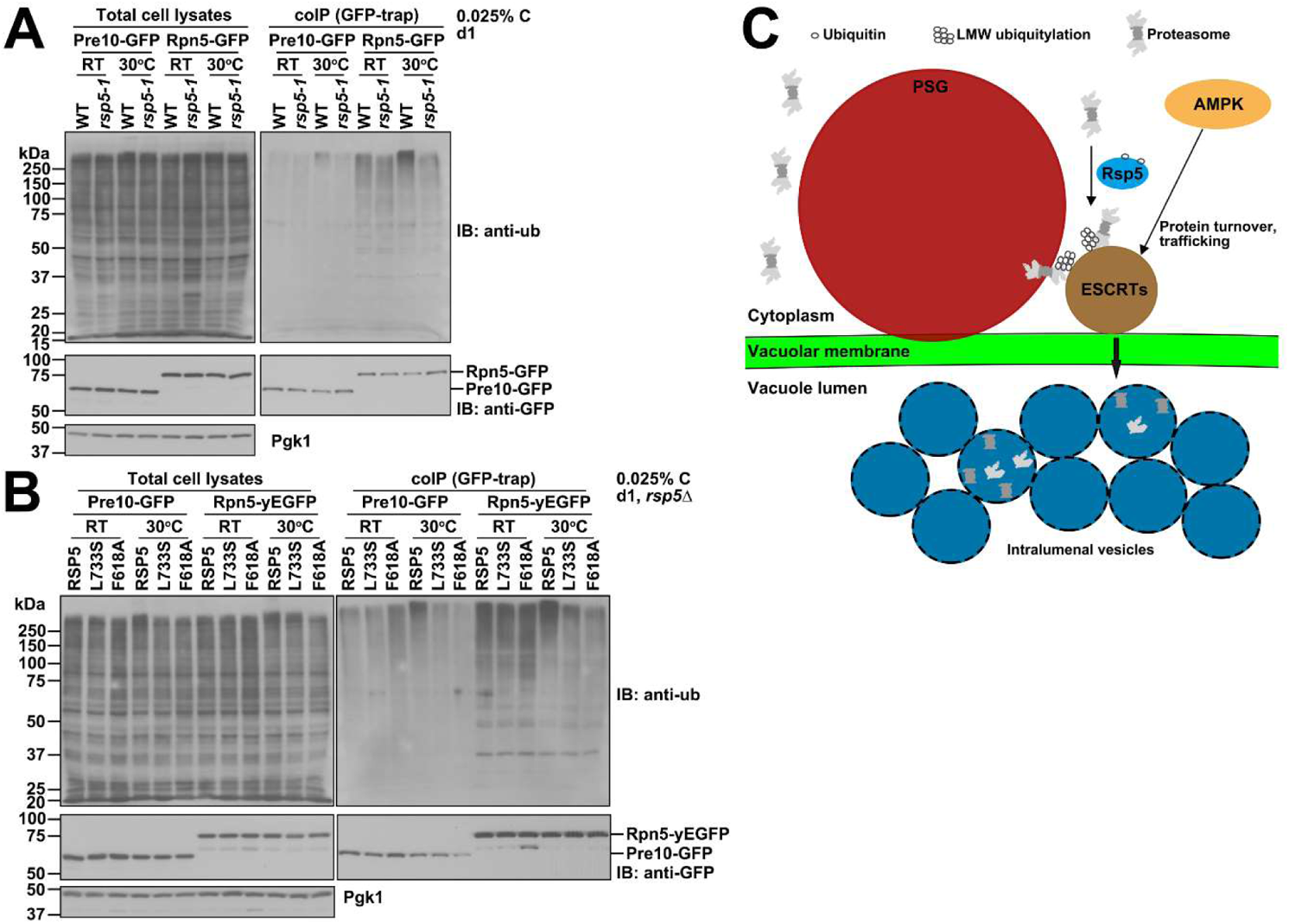
Rsp5 ubiquitin ligase is involved in proteasome ubiquitylation. (A) Co-IP analyses of ubiquitylated proteasome species in WT and rsp5-1 mutant cells expressing Pre10-GFP or Rpn5-GFP. Cultures were grown for ~1 d at 30°C. (B) Co-IP analyses of ubiquitylated proteasome species in rsp5Δ cells containing the indicated RSP5 alleles in pRS316. Cells were harvested from cultures in low glucose medium after ~1 d at 30°C. Representative blots are shown. (C) Working model of AMPK-directed selective microautophagy of proteasomes under low glucose conditions. AMPK activation in response to low glucose stress promotes Vps27 vacuolar trafficking and degradation. Vps27 (ESCRT-0) in turn sorts aberrant proteasomes associated with LMW ubiquitylation to the vacuole surface to initiate microautophagy. Proteasome RP subunits are the main targets for Rsp5.

To verify the importance of Rsp5 for proteasome ubiquitylation, we made a set of Rsp5-expressing constructs on CEN plasmids carrying RSP5, rsp5-L733S, and rsp5-F618A to disable the HECT domain noncovalent ubiquitin-binding site (French et al., 2009) and placed them in an rsp5Δ yeast background. As expected, plasmid-borne rsp5-L733S caused defects in proteasome ubiquitylation comparable to those seen in rsp5-1 cells at 30°C (Fig. 7A, B). The rsp5-F618A mutant also displayed defects in proteasome ubiquitylation (Fig. 7B), indicating that noncovalent ubiquitin binding by the Rsp5 HECT domain is important for proteasome ubiquitylation. Together, these results implicate the Rsp5 ligase in proteasome ubiquitylation in cells subject to low glucose stress.

## Discussion

Previously, we had identified a novel proteasome degradation pathway in low glucose conditions involving AMPK-regulated ESCRT-dependent microautophagy (Li et al., 2019). In this study, we sought to understand more about AMPK regulation and the mechanistic role of ESCRTs in proteasome microautophagy. In low carbon conditions, we find that AMPK promotes Vps27 protein turnover and its trafficking to the vacuole (Fig. 1). Vps27 is essential for ILV generation in vacuoles during microautophagy (Fig. 6C, D). Genetic or pharmacological induction of aberrant or inactive proteasomes leads to increased LMW ubiquitylation of these particles in cells under low glucose stress. Vps27 (ESCRT-0) appears to play an essential role in recognizing proteasome ubiquitylation and eliminating aberrant proteasomes through microautophagy (Figs. 4-6). The Rsp5 ubiquitin ligase contributes to proteasome ubiquitylation in this proteasome quality control pathway (Fig. 7A, B). Our results are summarized in the working model in Fig. 7C.

Exactly how AMPK regulates ESCRT-dependent microautophagy has yet to be determined. Previous studies have shown that TORC1 (Target of Rapamycin Complex 1) phosphorylates and antagonizes the function of Vps27 in microautophagy (Hatakeyama et al., 2019), while AMPK negatively regulates TORC1 activity through phosphorylation and aggregation of its Kog1 subunit, which results in disassembly of the TORC1 complex during glucose starvation (Hughes Hallett et al., 2015). The AMPK and TORC1 signaling pathways have interlinked and generally antagonistic effects (González et al., 2020). For ESCRT-dependent microautophagy in low glucose conditions, AMPK may inhibit TORC1 and thus relieve the suppressive effects of TORC1 on this process. It is also possible that AMPK competes with TORC1 and functions directly on ESCRT components for microautophagy initiation or activation, which would be reminiscent of the role of AMPK in macroautophagy (Kim et al., 2011).

Ubiquitylation of proteasomes could fine-tune their function in various ways, such as protein-proteasome interaction, proteasome activity, and cellular localization (Kors et al., 2019). At least 12 yeast proteasome subunits are known to be ubiquitylated in vivo and over 19 ubiquitylation sites have been detected (Hirano et al., 2015; Starita et al., 2012; Swaney et al., 2013). Only a small set of growth conditions and methods for proteasome purification have been reported in these studies, so the real numbers are likely to be substantially higher. Previous studies on possible functions of proteasome ubiquitylation are limited as well. In human prostate cancer cells, ubiquitylation of the CP subunit α2 favors the binding of delta-aminolevulinic acid dehydratase (ALAD); ALAD binds in place of 19S RP and functions as an endogenous proteasome inhibitor (Schmitt et al., 2016). RP ubiquitylation can also regulate proteasome activity. For example, polyubiquitylation of Rpn13 interferes with proteasome binding and degradation of ubiquitin-conjugated substrates in human cell lines (Besche et al., 2014), while monoubiquitylation of Rpn10 can limit the capacity of the UIM of Rpn10 to interact with substrates (Isasa et al., 2010). In our study, it is intriguing that aberrant proteasomes are preferably subject to LMW ubiquitylation (Fig. 5A), which appears to serve as a sorting signal for ESCRT-dependent clearance by microautophagy. It is unknown whether aberrant proteasomes are targeted by ESCRT-0 after their incorporation into PSGs or by bypassing PSGs altogether.

Ubiquitylation also plays a key role in proteasome macroautophagy (Marshall et al., 2016). The exact nutrient stress conditions help determine the distinct routes of proteasome autophagy in cells, and different proteasome ubiquitylation patterns may allow the diversification of proteasome autophagy routes, including the selective removal of aberrant or inactive proteasomes. Nitrogen starvation normally triggers nonselective proteasome macroautophagy that does not appear to involve extensive proteasome ubiquitylation (Marshall et al., 2016). By contrast, low glucose levels can simultaneously induce proteasome macroautophagy and microautophagy, and the latter appears to favor the removal of aberrant or inactive proteasomes that are either directly modified by LMW ubiquitylation or tightly bound to proteins that are modified in this way.

Chemical inactivation of proteasomes in yeast cells grown under nonstarvation conditions triggers a selective proteasome macroautophagy pathway accompanied by extensive ubiquitylation of the inactivated proteasomes (Marshall et al., 2016). Although proteasome ubiquitylation is a common feature in determining the selectivity of proteasome microautophagy and macroautophagy, the molecular components involved in these two selective proteasome autophagy pathways are different. For instance, the Cue5 autophagy receptor and the Hsp42 chaperone are engaged in the recognition of inactivated proteasomes for selective macroautophagy (Marshall et al., 2016) but are dispensable for selective proteasome microautophagy (JL and MH, unpublished data). On the other hand, the ESCRT-0 complex is involved in the recognition of aberrant proteasomes accumulated under low glucose conditions; its role in selective proteasome macroautophagy has not been examined. Insofar as ESCRTs are required in nonselective proteasome macroautophagy upon nitrogen starvation, we would predict ESCRTs are also involved in selective proteasome macroautophagy but likely in a different way than in microautophagy, such as autophagosome closure (Zhou et al., 2019).

Multiple components of the ESCRT-0, -I, and –II complexes contain UBDs, which provides structural plasticity for a wide variety of ubiquitylated cargo recognition and sorting mechanisms (Schmidt and Teis, 2012; Shields et al., 2009; Shields and Piper, 2011). In addition, having multiple UBDs can boost the avidity for ubiquitylated cargos as individual ESCRT UBDs have low affinity for ubiquitin; this might also allow recognition of very large cargo protein complexes with complicated ubiquitylation patterns (Piper et al., 2014), such as the LMW ubiquitylation of proteasomes we report here (Figs. 4, 5).

We still detected Vps27-proteasome RP interaction when ubiquitin binding by Vps27 or ESCRT-0 should have been eliminated, as in the vps27-VHS-UIM-PTAPL mutant (Fig. 5B-E). This suggests that ESCRT-0 can also interact with proteasomes in ways that are independent of RP ubiquitylation. It is possible that alternative binding partners may participate in these ESCRT-0-proteasome interactions. Nevertheless, the vps27-VHS-UIM-PTAPL mutation blocked ILV formation in the vacuole lumen to a degree comparable to the deletion mutant of VPS27 (Fig. 6). This implies that ubiquitin binding by ESCRT-0 is essential for microautophagy under carbon-limiting conditions. We note that the vps27-PTAPL mutant, which is defective for ESCRT-I and ESCRT-II binding, did not cause any major defects in proteasome degradation (Figs 5E, 6A), suggesting ESCRT-0 has additional sites for recruiting these downstream complexes, in line with previous findings (Shields and Piper, 2011).

In conclusion, we provide evidence here that ESCRT-0 functions in recognizing aberrant proteasomes through a proteasome ubiquitylation signature that makes these proteasomes favored targets for microautophagic elimination under low glucose stress, thereby ensuring that normal proteasomes, sequestered in PSGs, are readily available once a new source of glucose has been secured to allow a return to growth.

## Materials and Methods

### Yeast strains and cell growth

Yeast genetic manipulations were conducted using standard protocols (Dunham et al., 2015). Yeast strains used in this study are listed in Table S2. Unless specified in the figure legends, yeast cells were grown overnight at 30°C with vigorous agitation in synthetic complete (SC) medium (0.67% yeast nitrogen base without amino acids, 0.5% casamino acids, 0.002% adenine, 0.004% tryptophan, 0.002% uracil, and 2% glucose); tryptophan and uracil single or double dropout media were used for plasmid selection. Cells were then back-diluted in fresh SC medium and grown to mid-exponential phase. The cells were pelleted and rinsed once with sterile water, followed by different treatments. For glucose starvation, cells were resuspended in SC medium containing 0.025% glucose or lacking glucose and cultured for ~1 day or/and ~4 days at 30°C, as specified in the figures. For experiments with the proteasome inhibitor MG132, all strains included the pdr5Δ allele to allow efficient intracellular accumulation of the drug (Collins et al., 2010). Cells were grown in SC medium as above, and mid-log cells were resuspended in SC medium containing 0.025% glucose and DMSO or 50 μM MG132 (Santa Cruz Biotechnology, catalog # sc-201270, lot # H1219) dissolved in DMSO and cultured for ~1 day at 30°C.

### Plasmid construction

Genes of interest were amplified from yeast genomic DNA and cloned into the following plasmids: pRS316 (Sikorski and Hieter, 1989), p424GPD, or p426GPD (Mumberg et al., 1995) through restriction enzyme digestion and T4 DNA ligase-mediated ligation (NEB). The resulting plasmids were verified by enzymatic digestion and DNA sequencing. Quikchange site-directed mutagenesis (Agilent) was performed to generate the doa4-C571S and rsp5-L733S and rsp5-F618A mutants. Plasmids used in this study are listed in Table S3.

### Fluorescence microscopy

For epifluorescence microscopy, yeast cells were visualized on an Axioskop microscope (Carl Zeiss) equipped with a Plan-Apochromat 100×/1.40 oil DIC objective lens, a CCD camera (AxioCam MRm; Carl Zeiss), and a HBO100W/2 light source. Images were taken using AxioVision software with an auto exposure setup and processed using Adobe Photoshop CC software.

For time-lapse video recordings, yeast cells were viewed on an LSM 880 Airyscan NLO/FCS confocal microscope with an Alpha Plan-Apochromat 100×/1.46 NA oil objective lens. Excitation was performed with an argon laser at 488 nm, and emission was collected in the range of 493-556 nm for GFP imaging. Time-lapse images were acquired using Airyscan 32-channel ultra-sensitive area detector and the video was processed with ZEN software.

### Immunogold labeling electron microscopy

Yeast cells were fixed with 4% paraformaldehyde (PFA) and 0.2% glutaraldehyde in PBS for 30 min at 30°C followed by further fixation in 4% PFA for 1 h at 4°C. The fixed cells were rinsed once with PBS and resuspended in 10% gelatin. The solidified blocks were trimmed and placed in 2.3 M sucrose on a rotor overnight at 4°C, and then transferred to aluminum pins and frozen rapidly in liquid nitrogen. The frozen blocks were cut into 60 nm thick sections on a Leica Cryo-EM UC6 UltraCut using the Tokuyasu method (Tokuyasu, 1973). The cell sections were placed on carbon/Formvar-coated grids and floated in a dish of PBS for immunolabeling. Grids were placed section side down on drops of 0.1 M ammonium chloride to quench untreated aldehyde groups, then blocked for nonspecific binding on 1% fish skin gelatin in PBS. Single labeled grids were incubated with a primary rabbit anti-20S antibody (Enzo Life Sciences, catalog # BML-PW9355, lot # 05101719) at 1:200 dilution, and Protein A-gold beads (10 nm; Utrecht Medical Center) were used for secondary staining. The grids were rinsed in PBS, fixed with 1% glutaraldehyde for 5 min, rinsed again and stained with a mixture of 0.5% uranyl acetate to methylcellulose. Grids were viewed under a transmission electron microscope (FEI Tecnai G2 Spirit BioTWIN) at 80 kV. Images were taken using a SIS Morada 11-megapixel CCD camera and iTEM (Olympus) software. Acquired images were processed using Adobe Photoshop CC software.

### Protein extraction and Western blotting

Total proteins were extracted using the method of Kushnirov (2000), and Western blotting was performed as described previously with minor modifications (Li et al., 2016). The equivalent of one OD_600_ of cells were pelleted and washed once with sterile water. Cells were incubated in 400 μl 0.1 M NaOH for 5 min at room temperature (RT) and were then pelleted and resuspended in 100 μl SDS sample buffer (10% glycerol, 2% SDS, 0.1 M DTT, 62.5 mM Tris-HCl pH 6.8, 4% 2-mercaptoethanol, 0.008% bromophenol blue) and heated at 100°C for 5 min. Cell debris was removed by centrifugation. Equal volumes of the supernatants were loaded onto 10% (v/v) SDS-PAGE gels, followed by the transfer of proteins to polyvinylidene difluoride (PVDF) membranes (EMD Millipore, catalog # IPVH00010, lot # R0HB88840).

The membranes were incubated with the following primary antibodies: rabbit anti-Vps27 (Hatakeyama and De Virgilio, 2019) at 1:1000 dilution; JL-8 anti-GFP monoclonal antibody (TaKaRa, catalog # 632381, lot # A8034133) at 1:2,000 dilution; or an anti-Pgk1 monoclonal antibody (Abcam, catalog # ab113687, lot # GR3373682-5) at 1:10,000 dilution. Primary antibody binding was followed by anti-mouse (GE Healthcare, catalog # NXA931V, lot # 17041890) or anti-rabbit IgG (GE Healthcare, catalog # NA934V, lot # 17136627) secondary antibody conjugated to horseradish peroxidase at 1:10,000 dilution. The membranes were incubated in ECL detection reagent (Mruk and Cheng, 2011), and the ECL signals were detected using film (Thomas Scientific, catalog # 1141J52).

### Immunoprecipitation and proteasome ubiquitylation analysis

Cells equivalent to 35 OD_600_ units were pelleted and washed once with ice-cold sterile water. Cells were resuspended in 450 μl lysis buffer (20 mM Tris pH 7.5, 0.5 mM EDTA pH 8, 200 mM NaCl, 10% glycerol, 1 mM PMSF, 10 mM NEM, Roche cOmplete protease inhibitor catalog # 11836153001). Total cell lysates were prepared by beating with acid-washed glass beads using a FastPrep-24 device at 4°C for five cycles of 30 s agitation with 1 min breaks. After addition of 600 μl lysis buffer and Triton X-100 to 0.1% final concentration in cell lysates, cellular membranes were solubilized by incubating for 30 min on ice. Crude cell lysates were cleared at 16,000 x g for 10 min at 4°C. Cleared cell lysates were incubated with 25 μl GFP-Trap agarose resin slurry (ChromoTek) for 1 h to immunoprecipitate proteasomes. The resin was washed three times with lysis buffer containing 0.02% Triton X-100 (except that NaCl concentration was increased to 250 mM in Figure S4C) and resuspended in 50 μl 2 x SDS sample buffer. Bound proteins were eluted by incubating the beads for 10 min at 42°C. The elutes were analyzed by Western blotting as above with the following primary antibodies: polyclonal rabbit anti-ubiquitin (Dako, catalog # Z0458, lot # 20016967) at 1:2,000 dilution; monoclonal rabbit anti-ubiquitin-K63, clone Apu3 (EMD Millipore Corp, catalog # 05-1308, lot # 3137755) at 1:1,000 dilution; or monoclonal anti-HA (Sigma, catalog # H9658, lot # 089M4796V) at 1:2,000 dilution.

### Affinity purification of proteasomes

Proteasomes were affinity purified from yeast cells as described previously, with minor modifications (Li et al., 2015). Briefly, cells were harvested after growth under indicated conditions and grounded to fine powder with liquid nitrogen. About 6-7 ml of cell powders were thawed on ice, resuspended in 10 ml buffer A (50 mM Tris pH 7.5, 150 mM NaCl, 10% glycerol, 5 mM MgCl_2_, 5 mM ATP, Roche cOmplete EDTA-free protease inhibitor catalog # 11872580001), and incubated for 15 min on ice. Cell debris were pelleted at 30,000 x g for 20 min at 4°C. Total protein concentrations of the supernatants were determined using a Pierce BCA protein assay kit (Thermo Scientific, catalog # 23225, lot # SJ256254) according to the manufacturer’s protocol. Supernatant equivalent to ~100 mg total protein was incubated with 200 μl (packed) resin of anti-FLAG M2 affinity gel (Sigma, catalog # A2220, lot # SLCH0130) for 2 h on a rotator at 4°C. The proteasome-bound resin was washed twice with 12 ml buffer A for 10 min, and then incubated with 3 resin volumes of 200 µg·ml^-1^ 3xFLAG peptide (Sigma, catalog # F4799, lot # SLCJ4916) for 45 min to elute proteasome complexes. Proteasomes were concentrated with 100K MWCO centrifugal filters (Merk Millipore, catalog # UFC510024, lot # R9HA55989) and quantified with a BSA standard using a G:Box Chemi HR16 imager (Syngene).

### Protein digestion for LC-MS/MS

Protein samples were prepared as described above in the Immunoprecipitation section and resolved on 10% SDS-PAGE gels. Gel bands were excised and fixed with two incubations in 45% methanol and 10% acetic acid for 30 min at RT, and rinsed once with Milli Q water before the in-gel digestion procedure.

Gel bands were cut into small pieces and rinsed with 800 µl water on a tilt-table for 10 min and then washed for 20 min with 800 µl 50% acetonitrile (ACN)/100 mM NH_4_HCO_3_ (ammonium bicarbonate, ABC). The samples were reduced by incubating with 200 µl 4.5 mM DTT in 100 mM ABC for 20 min at 37°C, then alkylated by incubating with 200 µl 10 mM iodoacetamide in 100 mM ABC for 20 min at RT in the dark. The gels were washed for 20 min with 800 µl 50% ACN/100 mM ABC, then washed for 20 min with 800 µl 50% ACN/25 mM ABC. The gel pieces were briefly dried by SpeedVac, then resuspended in 200 µl of 25 mM ABC containing 500 ng of sequencing grade trypsin (Promega, catalog # V5111) and incubated for 16 h at 37°C. Peptides in the supernatant were moved to a new Eppendorf tube, and residual peptides were extracted from the gel by the addition of 500 µl 80% ACN/0.1% trifluoroacetic acid (TFA) and combined with the first supernatant. Peptides were dried in a SpeedVac and dissolved in 25 µl MS loading buffer (2% ACN, 1% TFA), with 5 µl injected for liquid chromatography-tandem mass spectrometry (LC-MS/MS) analysis.

### LC-MS/MS analysis

LC-MS/MS analysis was performed on a Thermo Scientific Q Exactive Plus equipped with a Waters nanoAcquity UPLC system utilizing a binary solvent system (A: 100% water, 0.1% formic acid; B: 100% acetonitrile, 0.1% formic acid). Trapping was performed at 5 µl·min^-1^, 99.5% Buffer A for 3 min using a Waters ACQUITY UPLC M-Class Symmetry C18 Trap Column (100 Å, 5 µm, 180 µm x 20 mm, 2G, V/M). Peptides were separated at 37°C using a Waters ACQUITY UPLC M-Class Peptide BEH C18 Column (130 Å, 1.7 µm, 75 µm X 250 mm) and eluted at 300 nl·min^-1^ with the following gradient: 3% buffer B at initial conditions; 5% B at 1 min; 25% B at 90 min; 50% B at 110 min; 90% B at 115 min; 90% B at 120 min; return to initial conditions at 125 min. Mass spectra was acquired in profile mode over the 300-1,700 m/z range using 1 microscan, 70,000 resolution, AGC target of 3E6, and a maximum injection time of 45 ms. Data-dependent MS/MS were acquired in centroid mode on the top 20 precursors per MS scan using 1 microscan, 17,500 resolution, AGC target of 1E5, maximum injection time of 100 ms, and an isolation window of 1.7 m/z. Precursors were fragmented by HCD activation with a collision energy of 28%. MS/MS were collected on species with an intensity threshold of 1E4, charge states 2-6, and peptide match preferred. Dynamic exclusion was set to 20 sec.

### Peptide identification

Tandem mass spectra were extracted by Proteome Discoverer software (version 2.2.0.388, Thermo Scientific) and searched in-house using the Mascot algorithm (version 2.7.0, Matrix Science). The data were searched against the Swissprotein database with taxonomy restricted to Saccharomyces cerevisiae (7,905 sequences). Search parameters included trypsin digestion with up to 2 missed cleavages, peptide mass tolerance of 10 ppm, and MS/MS fragment tolerance of 0.02 Da. Cysteine carbamidomethylation was configured as a fixed modification. Methionine oxidation; phosphorylation of serine, threonine and tyrosine; propionamide adduct to cysteine; and GG dipeptide (ubiquitin residue) on lysine were configured as variable modifications. Normal and decoy database searches were run, with the confidence level was set to 95% (p<0.05). Scaffold Q+S (version 4.11.0, Proteome Software Inc.) was used to validate MS/MS based peptide and protein identifications. Peptide identifications were accepted if they could be established at greater than 95.0% probability by the Scaffold Local FDR algorithm. Protein identifications were accepted if they could be established at greater than 99.0% probability and contained at least 2 identified peptides.

### Statistical analysis

GraphPad Prism 9 was used to generate the heat map in Figure S6B and most bar graphs. Microsoft Excel 2013 was used to perform ANOVA single factor analysis and generate bar graphs in Figures 6D-6F. ImageJ was used to quantify protein band intensities in Figures 5B-5D. The number of cells counted for image quantification are described in the figure legends. Each experiment was repeated at least three times and the percentages shown in the figures represent the average of all the experiments. Error bars represent standard deviations.

## Acknowledgements

We thank Carolyn Breckel and Mengwen Zhang in the lab, and Xiaofeng Wang at Virginia Tech for critical reading of the manuscript. We thank Masahide Oku and Yasuyoshi Sakai at Kyoto University, and Riko Hatakeyama and Claudio De Virgilio at University of Fribourg for sharing antibodies, yeast strains, or plasmids; Morven Graham for electron microscopy assistance at the Yale University Center for Cellular and Molecular Imaging; and Jean Kanyo for LC-MS/MS assistance at the Yale University Keck MS & Proteomics Resource.

## Competing interests

The authors declare that no competing interests exist.

## Funding

Yale University Keck MS & Proteomics Resource was in part supported by National Institutes of Health grant SIG OD018034. This work was supported by National Institutes of Health grants GM136325 and GM046904 to M.H.

## Supplemental information

**Fig. S1.**
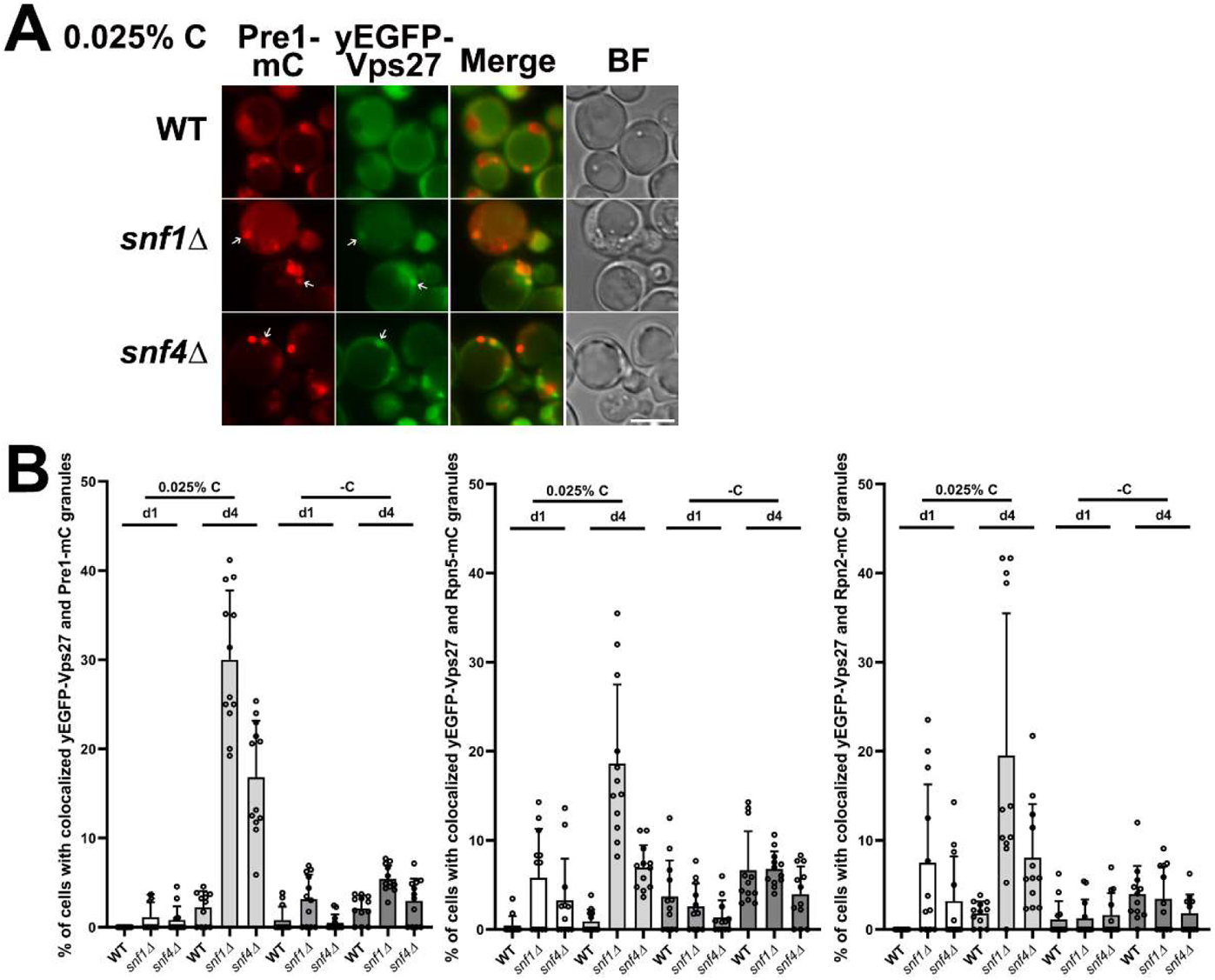
Vps27 is enriched in proteasome puncta in AMPK mutant cells under low glucose conditions. (A) Epifluorescence images of Pre1-mC (a CP subunit) in WT and AMPK mutant (snf1Δ or snf4Δ) cells expressing chromosomally integrated yEGFP-VPS27. Endogenous VPS27 was deleted. Cells were harvested from cultures grown in low glucose medium for ~4 d at 30°C. White arrows point to colocalized Vps27 and proteasome foci in AMPK mutant cells. BF, bright field. Scale bar, 5 µm. Representative images are shown. (B) Percentage of cells with colocalized yEGFP-Vps27 and proteasomal puncta (Pre1-mC, Rpn5-mC, or Rpn2-mC) in WT or AMPK mutant cells. Cultures were grow in low glucose or glucose-free medium for ~1 d and ~4 d at 30°C. Cells counted in WT: Pre1-mC (n=535, 444 for low glucose at day 1 and day 4, respectively, and n=551, 632 for glucose-free at day 1 and day 4, respectively); Rpn5-mC (560, 538, 446, 494); Rpn2-mC (337, 598, 426, 510) Cell counts in snf1Δ: Pre1-mC (194, 541, 517, 392); Rpn5-mC (194, 413, 538, 462), and Rpn2-mC (268, 381, 569, 368). Cell counts in snf4Δ: Pre1-mC (511, 622, 766, 364); Rpn5-mC (503, 432, 717, 322); and Rpn2-mC (488, 506, 669, 240). Results plotted as mean±sd..

**Fig. S2.**
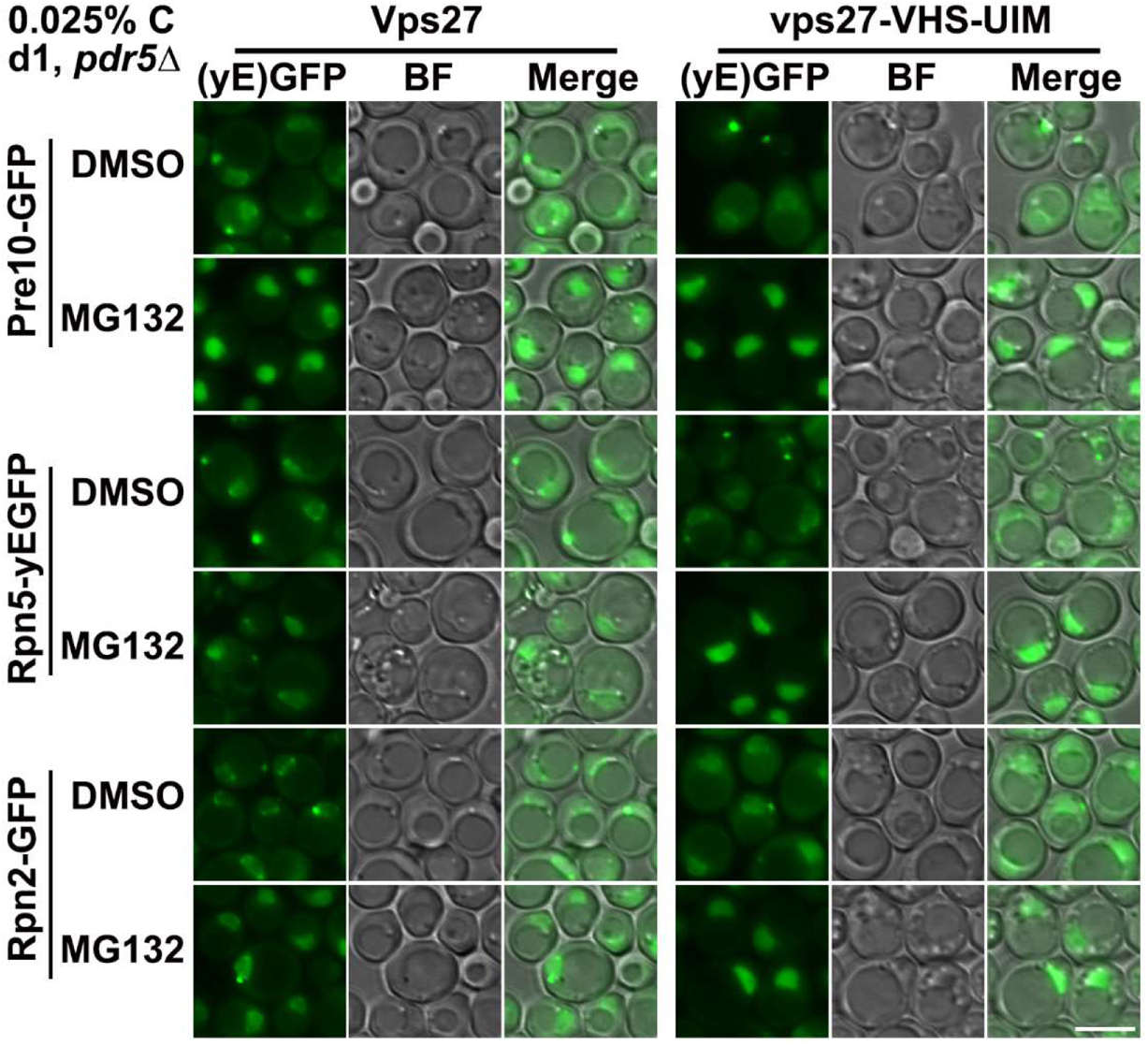
Effect of UBD mutations of Vps27 on inactive proteasome trafficking under low glucose conditions. Epifluorescence images of Pre10-GFP, Rpn5-yEGFP, and Rpn2-GFP in pdr5Δ cells expressing chromosomally integrated VPS27 and vps27-VHS-UIM. Endogenous VPS27 gene was deleted. Cultures were grown in low glucose medium containing DMSO or 50 µM MG132 for ~1 d at 30°C. The same cells were used for quantification as shown in panel Figure 2F. The vps27-VHS-UIM mutation further inhibited PSG formation in cells with MG132 treatment. BF, bright field. Scale bar, 5 µm. Representative images are shown.

**Fig. S3.**
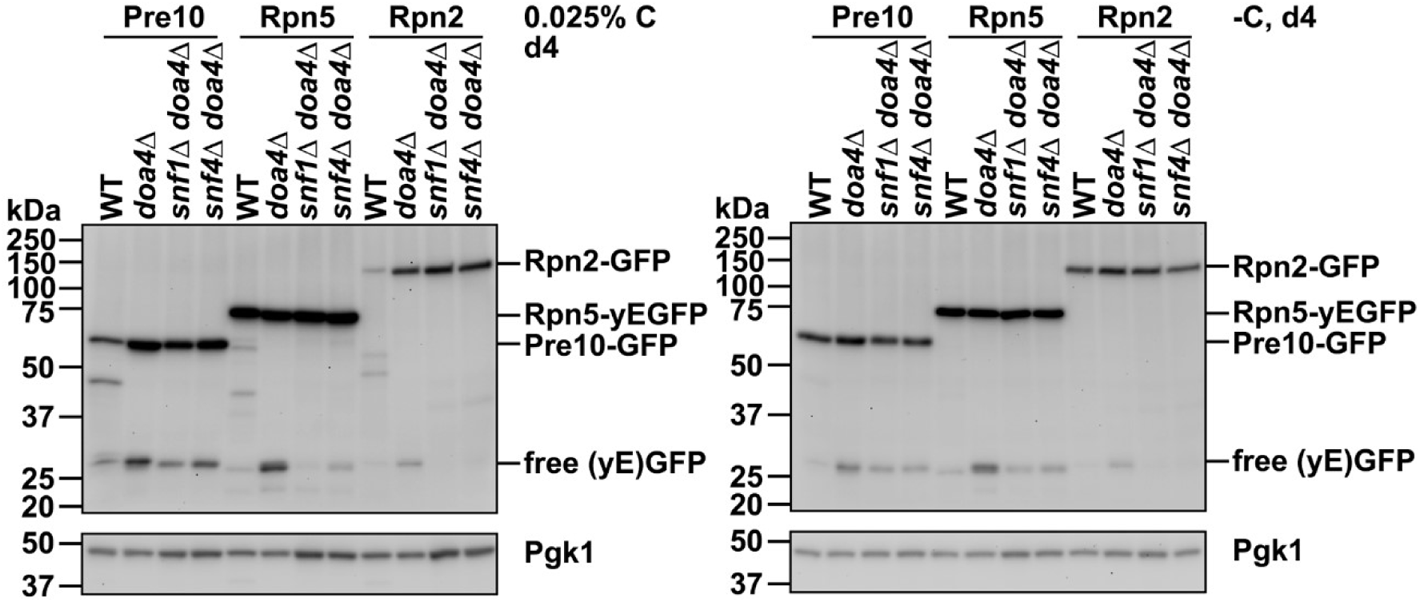
AMPK mutations do not suppress proteasome fragmentation defects in doa4Δ cells. Anti-GFP immunoblot analyses of free (yE)GFP release and proteasome fragmentation of Pre10-GFP, Rpn5-yEGFP, and Rpn2-GFP in the indicated strains. Cultures were grown in low glucose or glucose-free medium for ~4 d at 30°C. Representative blots are shown.

**Fig. S4.**
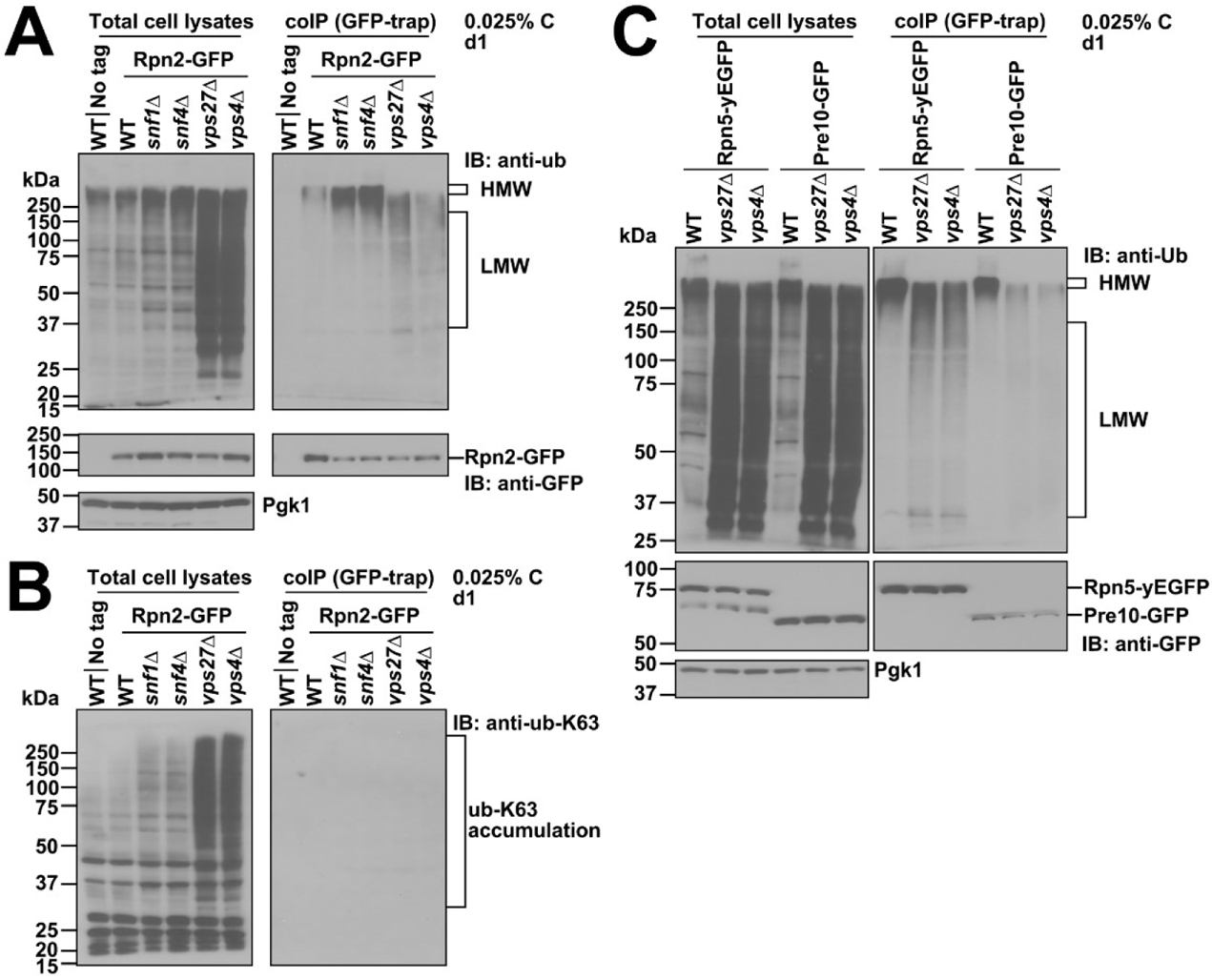
Ubiquitin-K63 chains are not detectably conjugated to proteasomes. (A) Co-IP analyses of ubiquitylated proteasome species in the indicated strains expressing Rpn2-GFP. LMW ubiquitylated proteasomal species accumulated in ESCRT mutants. (B) Anti-ubiquitin-K63 immunoblot analyses of the coIP samples from panel (A). The ubiquitin-K63-linked ubiquitylated species accumulated in total cell lysates of ESCRT mutants but were not detected in the isolated proteasomes. (C) Co-IP analyses of ubiquitylated proteasome species using GFP-trap agarose beads and washing in high salt (250 mM NaCl) buffer before elution. LMW ubiquitylated proteasome species were retained in ESCRT mutants. Cultures were in low glucose for ~1 d at 30°C.

**Fig. S5.**
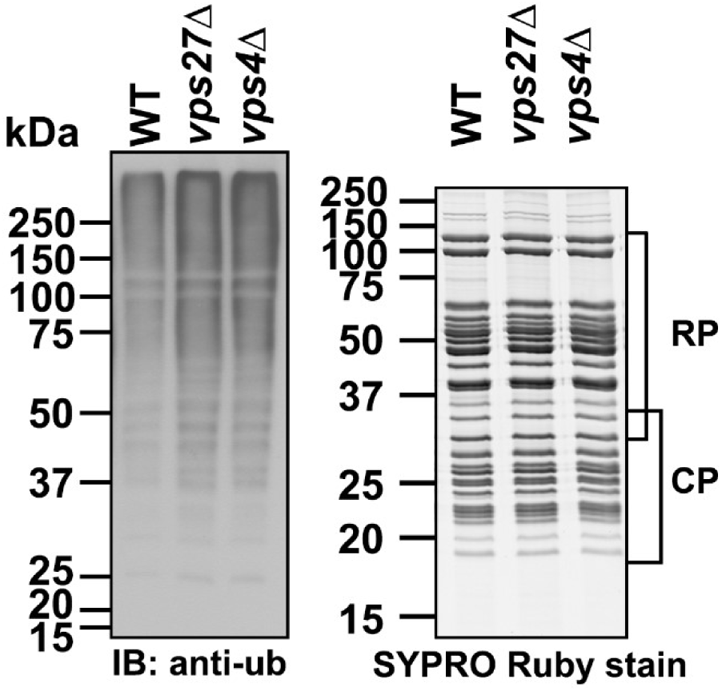
Ubiquitylated proteasome species accumulate in the purified proteasomes of ESCRT mutants under low glucose conditions. Anti-ubiquitin immunoblot analyses and SYPRO Ruby staining of purified proteasomes (10 µg) of WT and ESCRT mutant (vps27Δ, vps4Δ) cells. Cells were harvested from 700-ml cultures grown in low glucose medium for ~1 d at 30°C, and lysates were made by grinding cells in liquid nitrogen with ATP present in the buffer. Proteasomes were affinity purified through 3xFLAG tag fused on Rpn11 C-terminus and eluted with FLAG peptide.

**Fig. S6.**
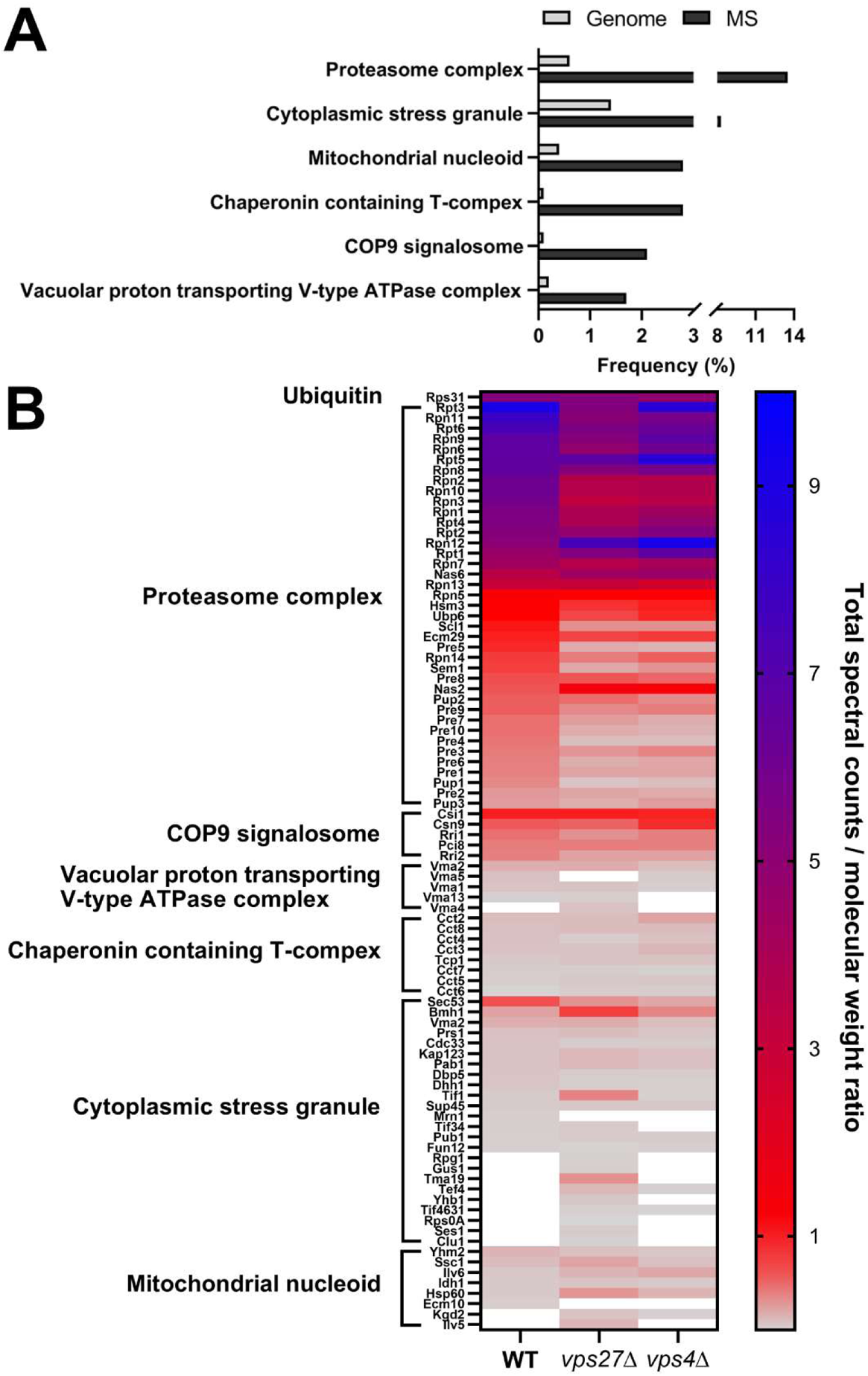
Ubiquitin, proteasome subunits and proteasome-associated proteins are dominant proteins seen by LC-MS/MS of the GFP-Trap elutes from WT and ESCRT mutant cells expressing Rpn5-yEGFP. (A) Gene ontology (GO) analysis of the protein hits from WT, vps27Δ, and vps4Δ cells, with cellular component ontology with P≤0.01. GO term analysis was conducted using the Generic Gene Ontology Term Finder online software (Princeton U.). (B) The ratio of total spectral counts to molecular weight of individual protein hits with enriched GO cellular component term as shown in panel (A). White color represents specific proteins that were not detected in the indicated strains. A full list of protein hits is given in Table S1.

**Fig. S7.**
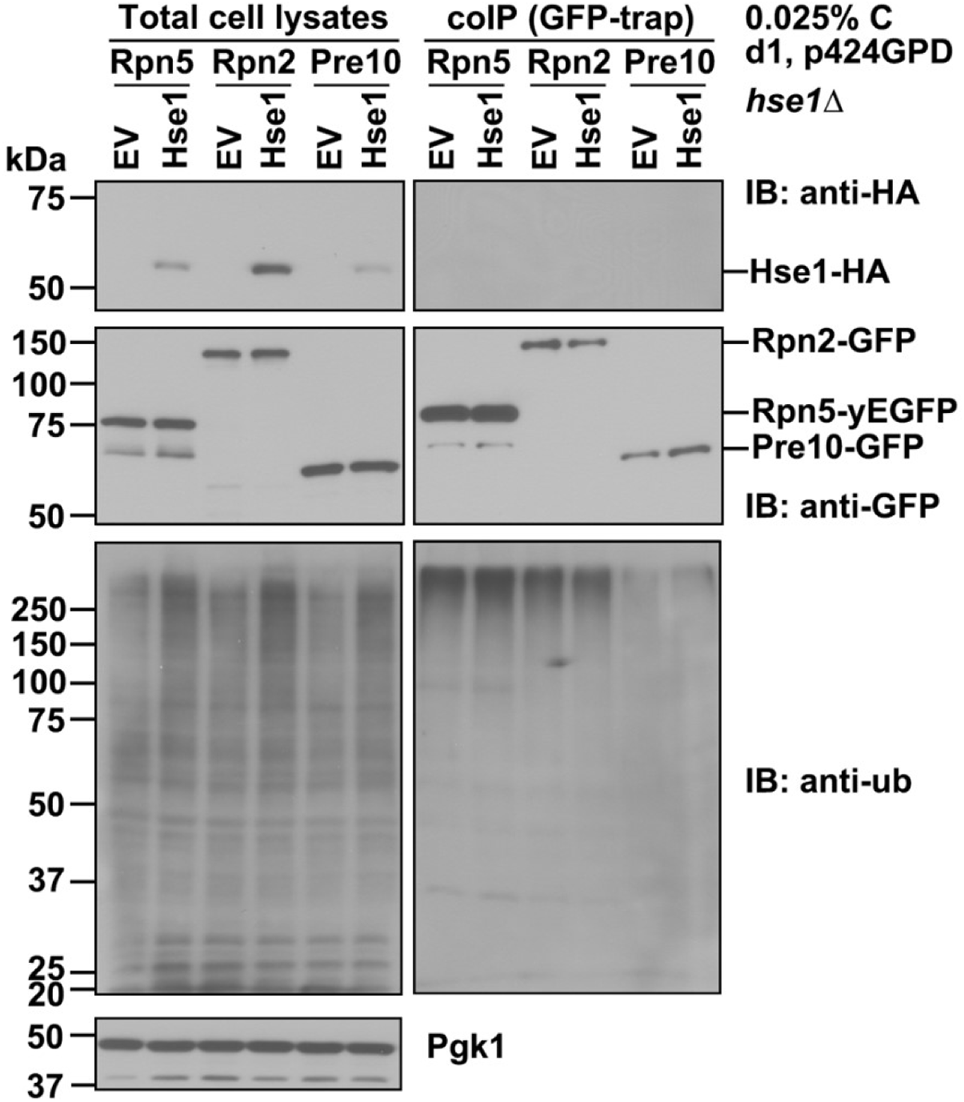
Overexpressed Hse1-HA by itself does not interact with proteasomes under low glucose conditions. Co-IP analyses of Hse1 and proteasomes in hse1Δ cells expressing Rpn5-yEGFP, Rpn2-GFP, and Pre10-GFP, as well as Hse1-HA under the GPD promoter with a p424GPD backbone. Proteasome ubiquitylation was not affected in hse1Δ cells. Cultures were grown in low glucose for ~1 d at 30°C.

**Fig. S8.**
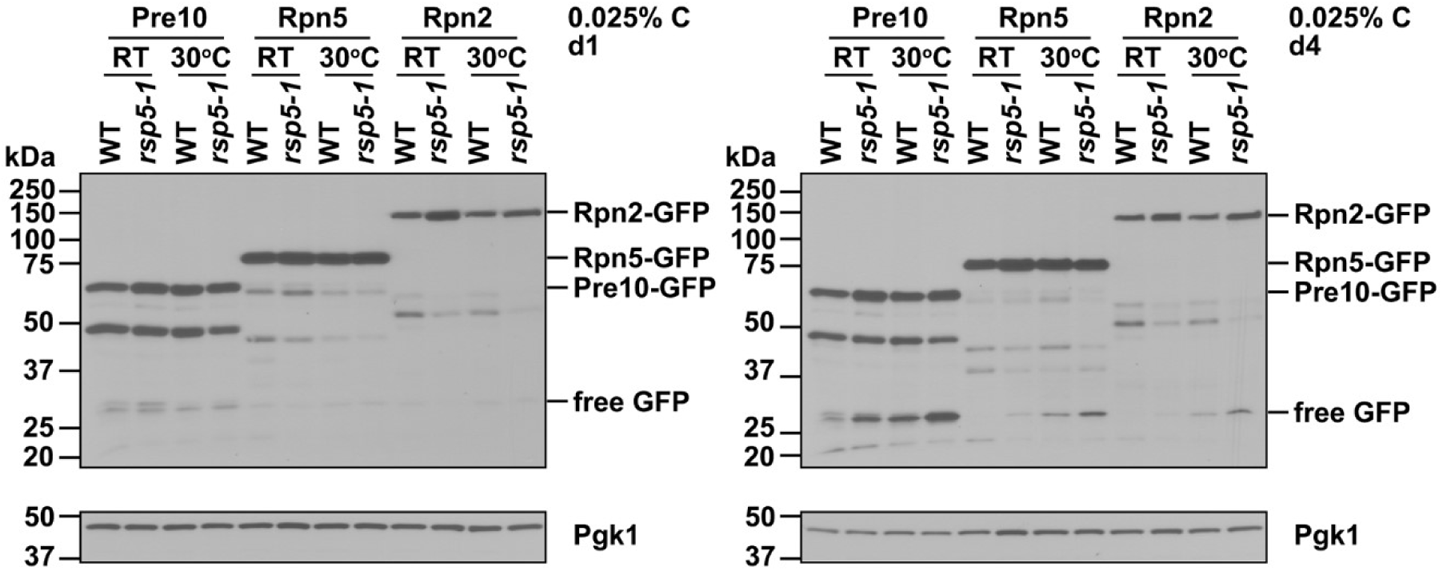
Proteasome RP subunit fragmentation is more sensitive to defective Rsp5 than that of a CP subunit. Anti-GFP immunoblot analyses of free GFP release and proteasome fragmentation of Pre10-GFP, Rpn5-GFP, and Rpn2-GFP in WT and rsp5-1 cells. Representative blots are shown.

## Supplementary tables

**Table S1 is in a separate Excel file**

**Table S1.** Proteins identified by mass spectrometry analyses of co-IP samples from WT and ESCRT mutant (vps27Δ, vps4Δ) cells expressing Rpn5-yEGFP. Note: Protein hits are sorted by total spectral counts of individual proteins in the WT sample from high to low. Peptide thresholds ≥ 95% and protein thresholds ≥ 99% and ≥ 2 peptides.

**Table S2.**
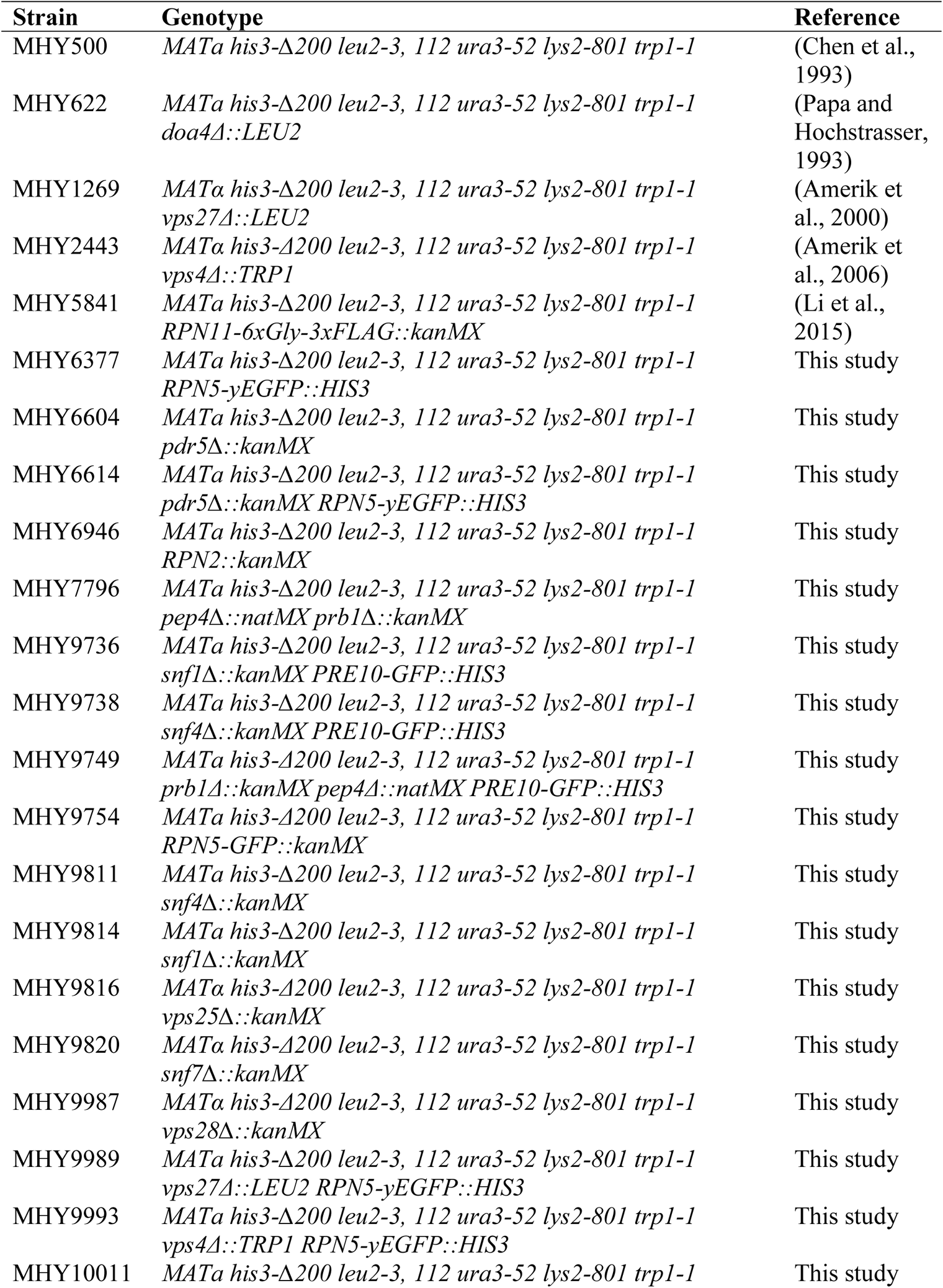
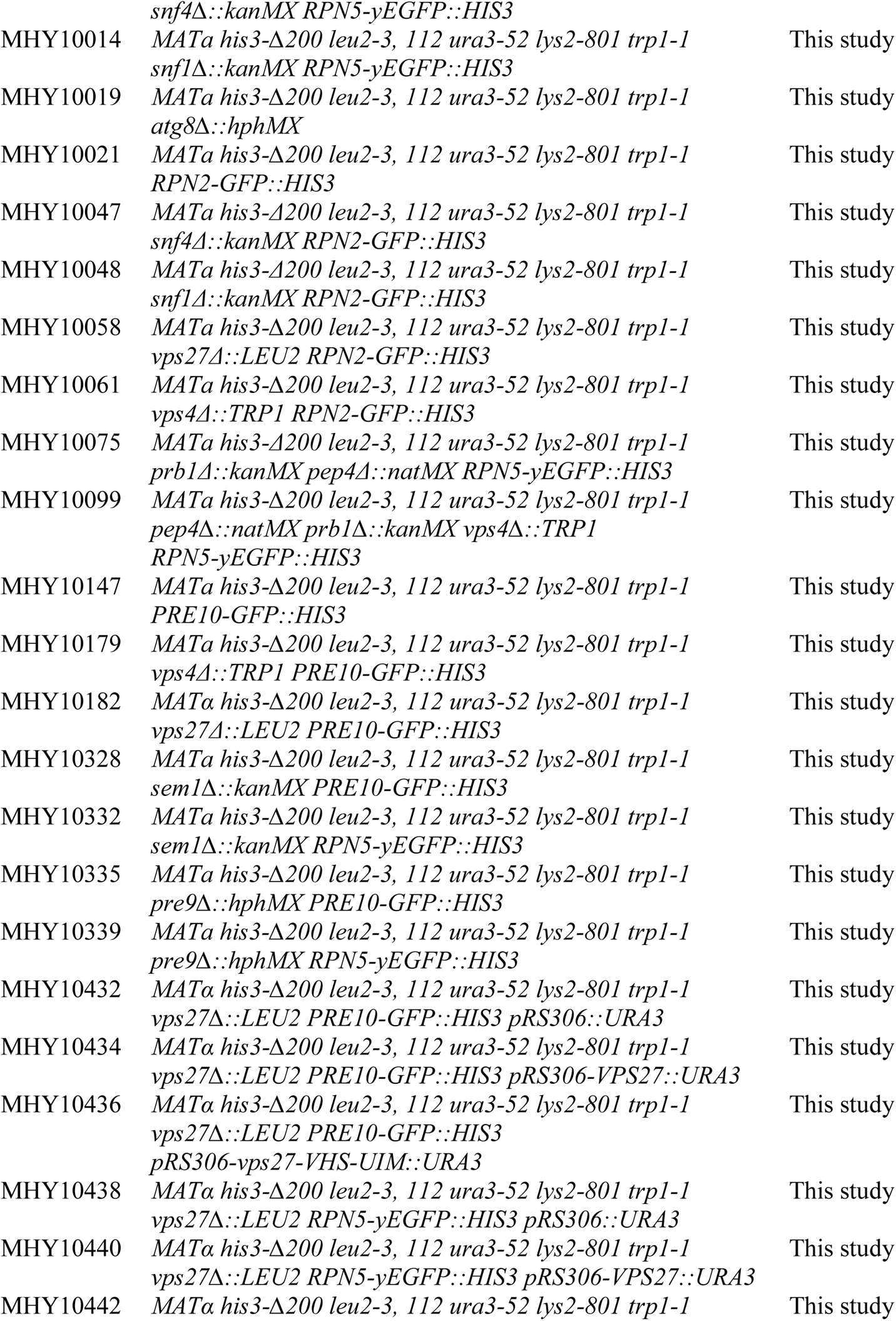
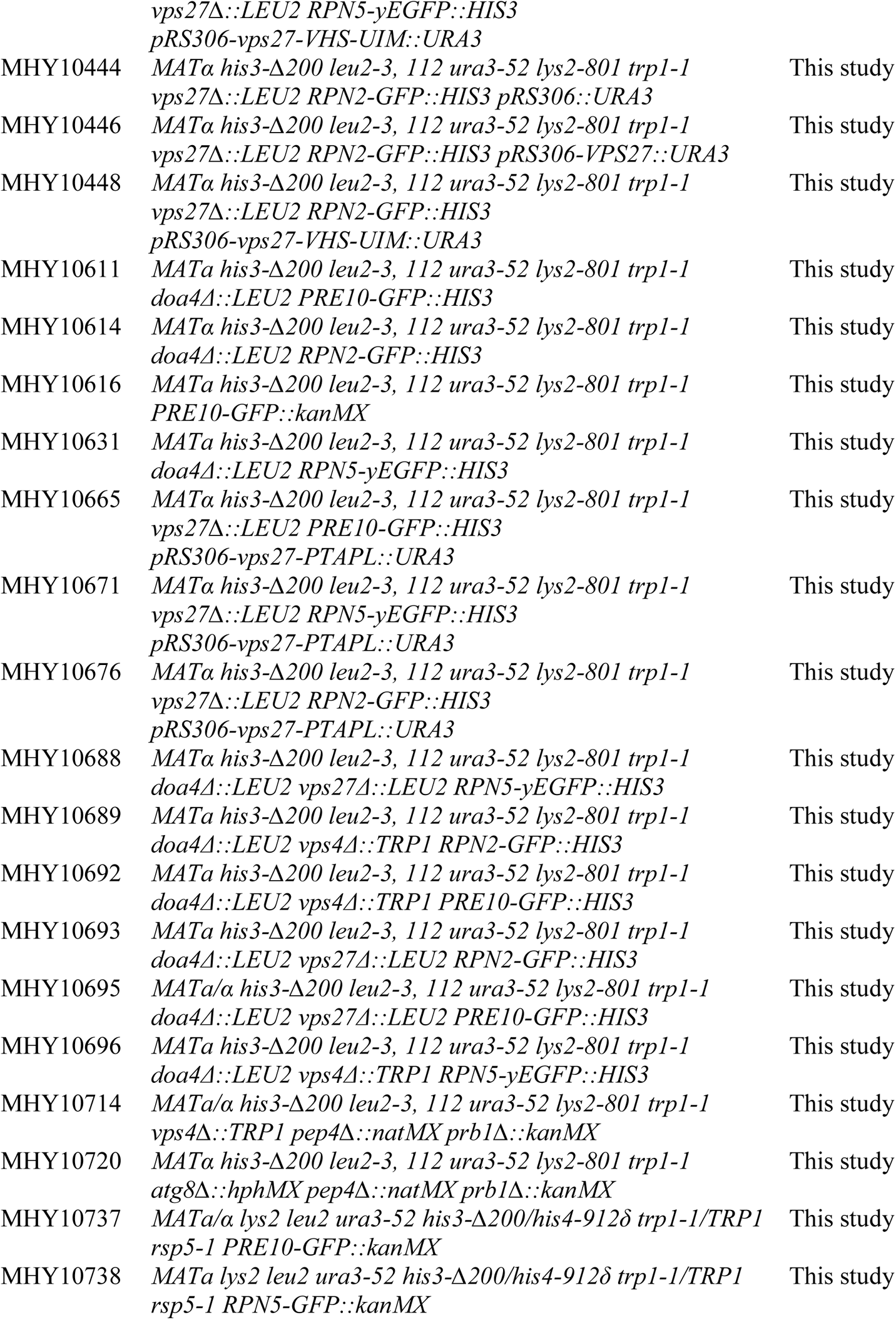
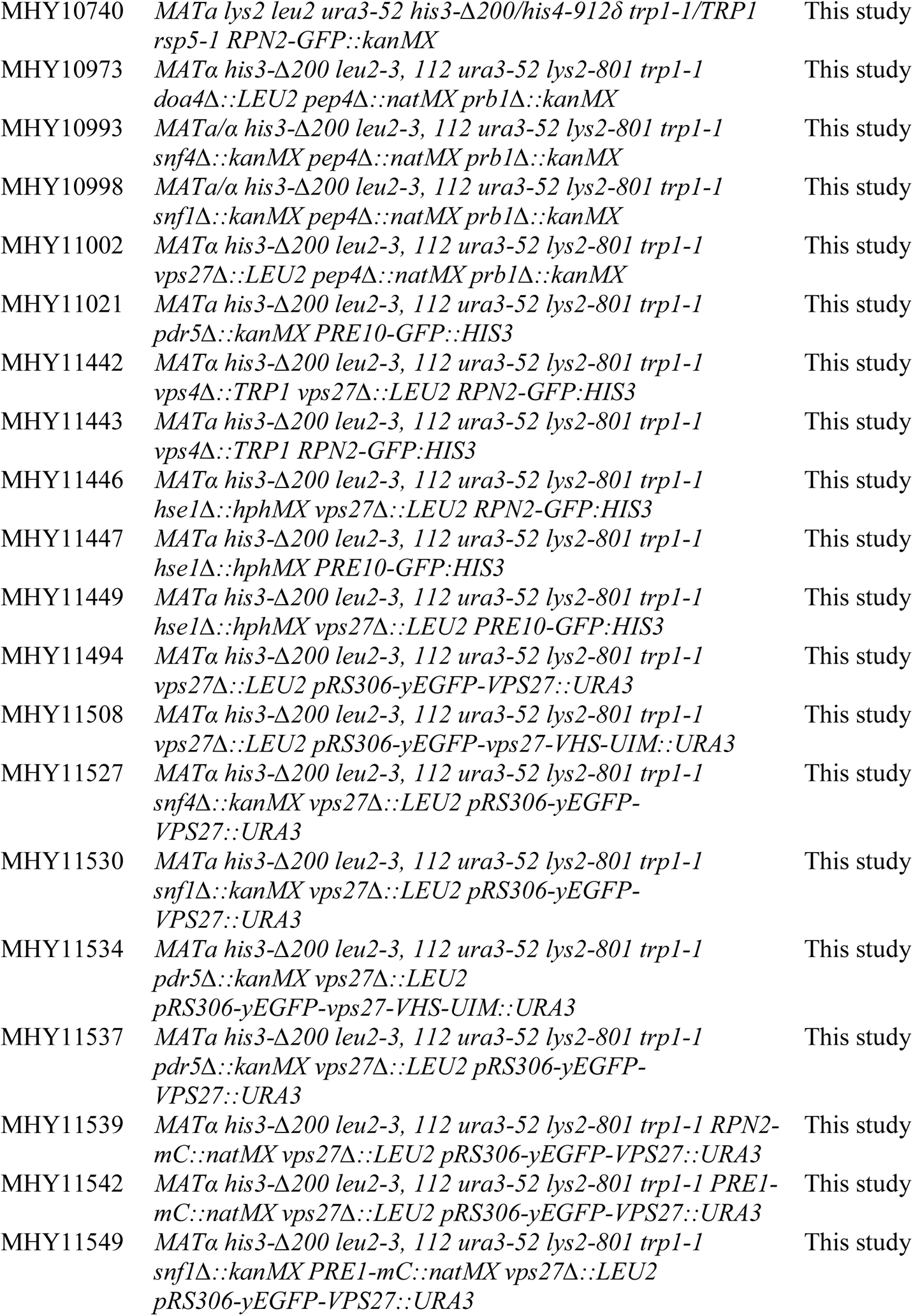
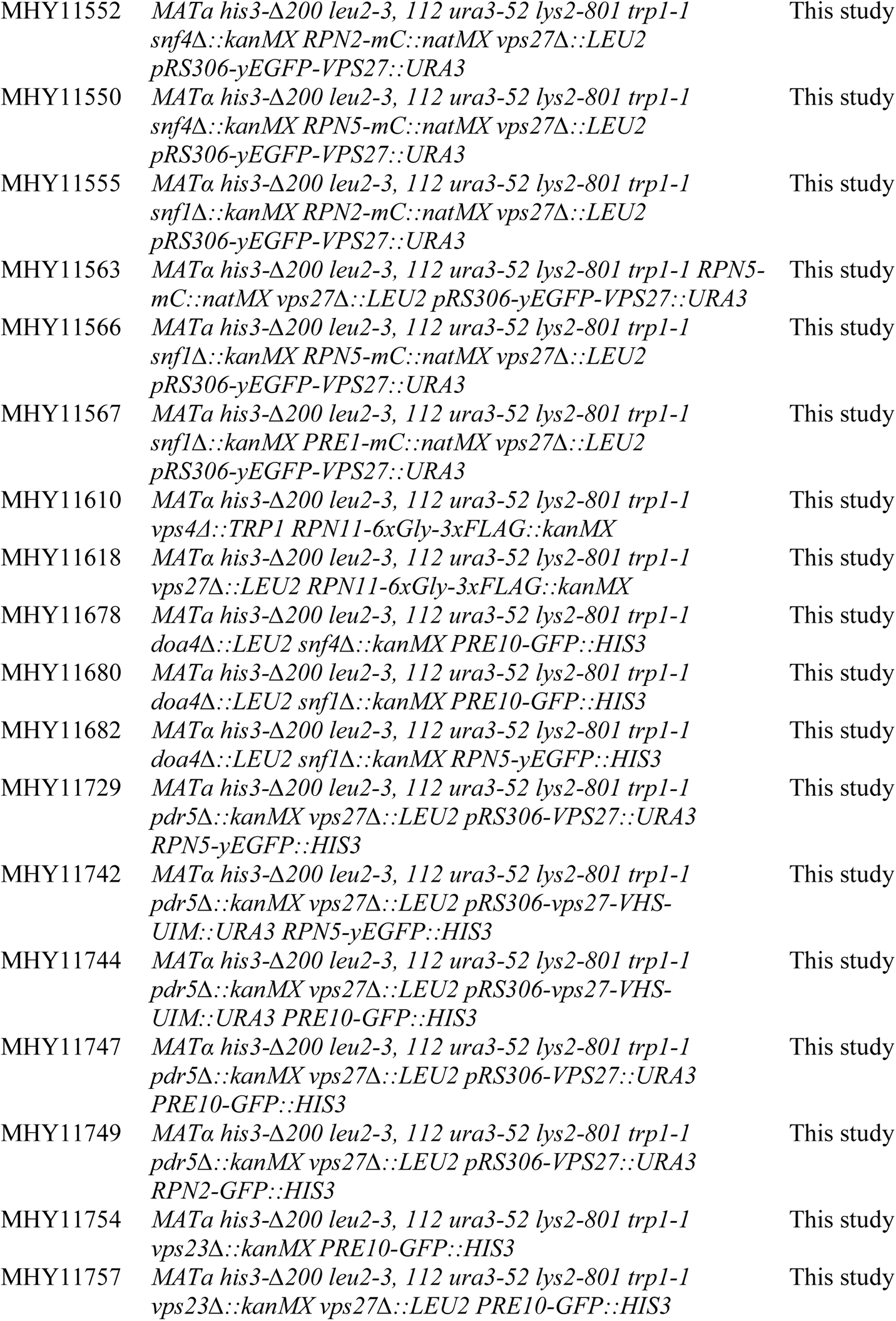
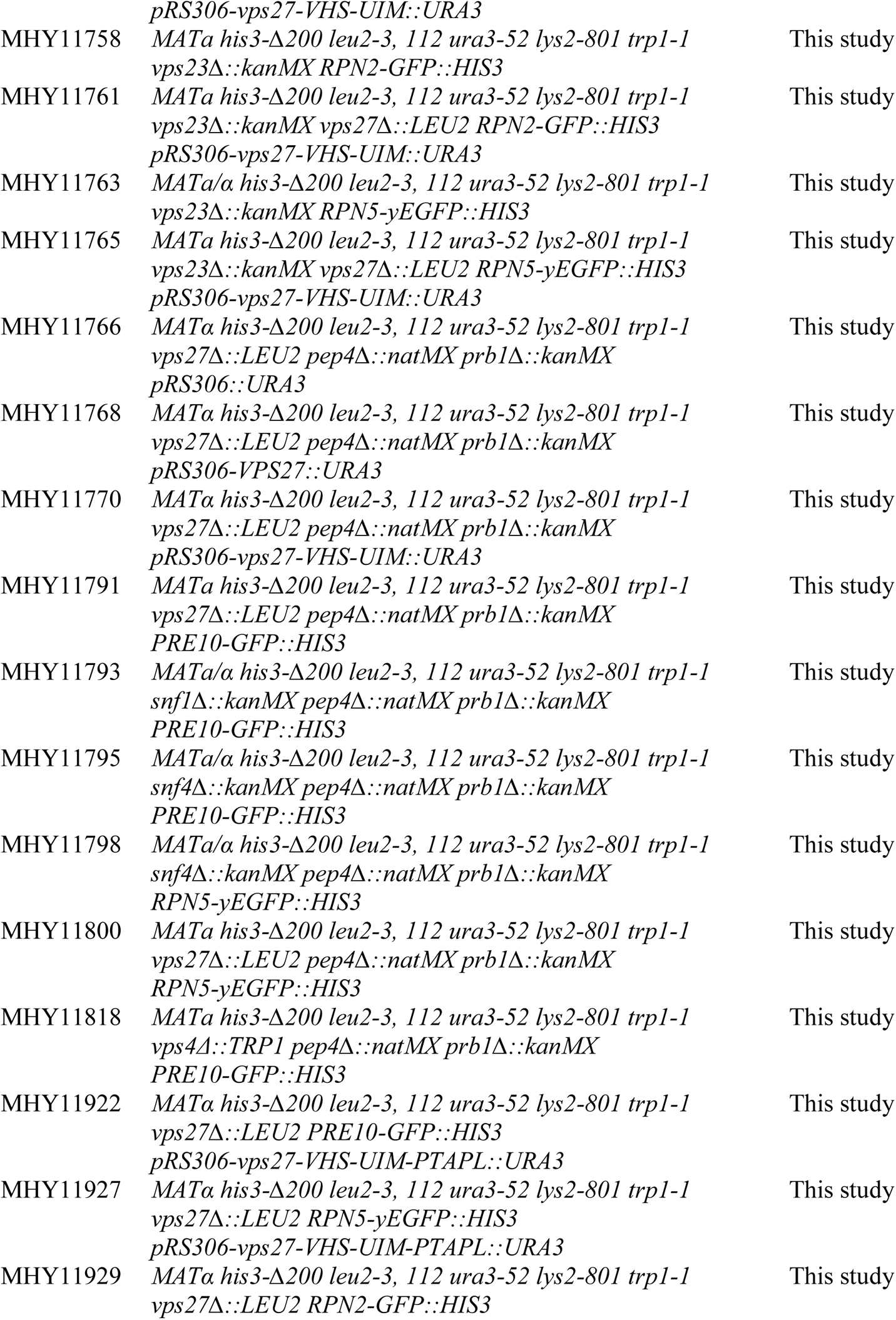
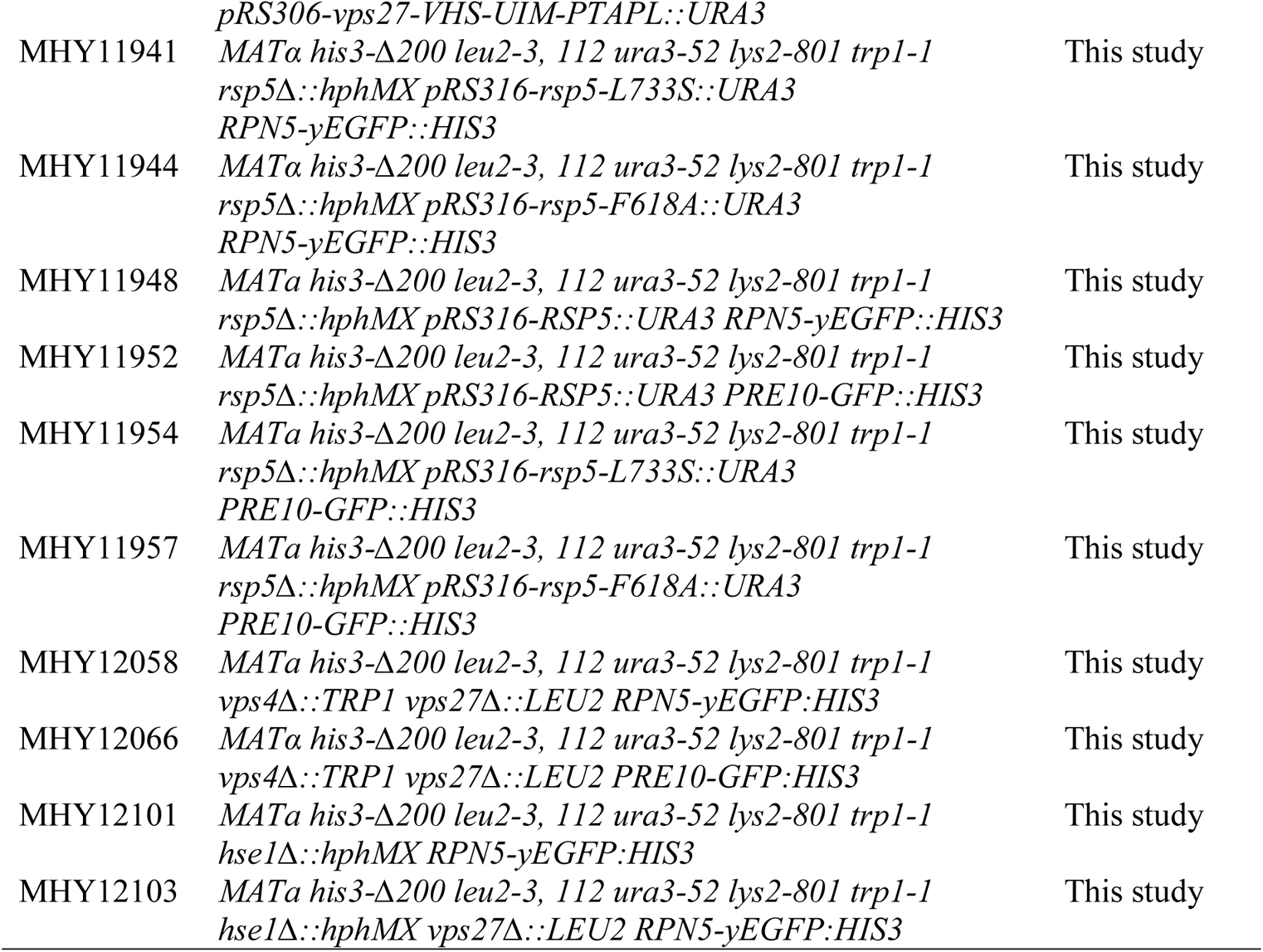
Yeast strains used in this study.

---

**Table S3.**
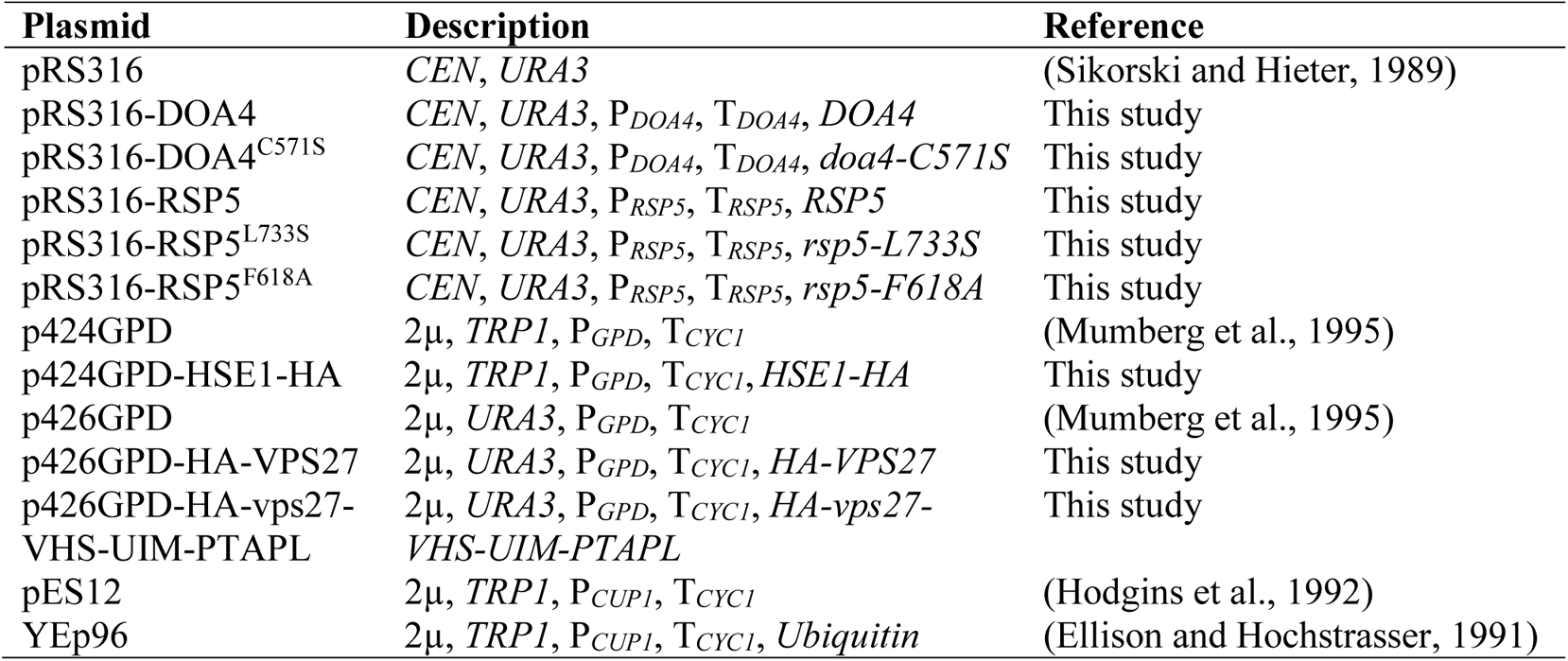
Plasmids used in this study.

**Movie 1 is in a separate Mov file**

**Movie 1. Defective vacuolar trafficking of yEGFP-Vps27 in snf1Δ cells under low glucose conditions.**

Confocal time-lapse images of yEGFP-Vps27 showing that perivacuolar granules of yEGFP-Vps27 accumulated in snf1Δ cells grown in low glucose medium for ~1 day at 30°C. The Vps27 granules were highly mobile, undergoing rapid docking and separation. The time-lapse video comprises 30 image frames taken at 5 s intervals with 2.53 s scanning time for each frame and played at 2 frames per second; the real time length was 145 s for this video.

## Notes

### Competing Interest Statement

The authors have declared no competing interest.

## References

Amerik, A. Y., Nowak, J., Swaminathan, S. and Hochstrasser, M. (2000). The Doa4 deubiquitinating enzyme is functionally linked to the vacuolar protein-sorting and endocytic pathways. Mol Biol Cell 11, 3365–3380.

Besche, H. C., Sha, Z., Kukushkin, N. V., Peth, A., Hock, E.-M., Kim, W., Gygi, S., Gutierrez, J. A., Liao, H., Dick, L. et al. (2014). Autoubiquitination of the 26S proteasome on Rpn13 regulates breakdown of ubiquitin conjugates. EMBO J 33, 1159–1176.

Bilodeau, P. S., Urbanowski, J. L., Winistorfer, S. C. and Piper, R. C. (2002). The Vps27p–Hse1p complex binds ubiquitin and mediates endosomal protein sorting. Nat Cell Biol 4, 534.

Bilodeau, P. S., Winistorfer, S. C., Kearney, W. R., Robertson, A. D. and Piper, R. C. (2003). Vps27-Hse1 and ESCRT-I complexes cooperate to increase efficiency of sorting ubiquitinated proteins at the endosome. J Cell Biol 163, 237–243.

Budenholzer, L., Cheng, C. L., Li, Y. and Hochstrasser, M. (2017). Proteasome structure and assembly. J Mol Biol 429, 3500–3524.

Carling, D. (2017). AMPK signalling in health and disease. Curr Opin Cell Biol 45, 31–37.

Chen, H. I. and Sudol, M. (1995). The WW domain of Yes-associated protein binds a proline-rich ligand that differs from the consensus established for Src homology 3-binding modules. Proc Natl Acad Sci USA 92, 7819–7823.

Coccetti, P., Nicastro, R. and Tripodi, F. (2018). Conventional and emerging roles of the energy sensor Snf1/AMPK in Saccharomyces cerevisiae. Microbial Cell 5, 482–494.

Collins, G. A., Gomez, T. A., Deshaies, R. J. and Tansey, W. P. (2010). Combined chemical and genetic approach to inhibit proteolysis by the proteasome. Yeast 27, 965–974.

Dunham, M. J., Gartenberg, M. R. and Brown, G. W. (2015). Methods in yeast genetics and genomics: Cold Spring Harbor Laboratory Press.

Enenkel, C. (2014). Proteasome dynamics. BBA-Mol Cell Res 1843, 39–46.

Finley, D., Ulrich, H. D., Sommer, T. and Kaiser, P. (2012). The ubiquitin– proteasome system of Saccharomyces cerevisiae. Genetics 192, 319–360.

French, M. E., Kretzmann, B. R. and Hicke, L. (2009). Regulation of the RSP5 ubiquitin ligase by an intrinsic ubiquitin-binding site. J Biol Chem 284, 12071–12079.

Gaullier, J.-M., Simonsen, A., D’Arrigo, A., Bremnes, B., Stenmark, H. and Aasland, R. (1998). FYVE fingers bind PtdIns(3)P. Nature 394, 432–433.

Ghillebert, R. (2011). The AMPK/SNF1/SnRK1 fuel gauge and energy regulator: structure, function and regulation Regulation of AMPK/SNF1/SnRK1 kinase complexes. FEBS J 278, 3978–3990.

González, A., Hall, M. N., Lin, S.-C. and Hardie, D. G. (2020). AMPK and TOR: The Yin and Yang of cellular nutrient sensing and growth control. Cell Metab 31, 472–492.

Hatakeyama, R. and De Virgilio, C. (2019). TORC1 specifically inhibits microautophagy through ESCRT-0. Curr Genet 65, 1243–1249.

Hatakeyama, R., Péli-Gulli, M.-P., Hu, Z., Jaquenoud, M., Garcia Osuna, G. M., Sardu, A., Dengjel, J. and De Virgilio, C. (2019). Spatially distinct pools of TORC1 balance protein homeostasis. Mol Cell 73, 325–338.e8.

Hecht, K. A., O’Donnell, A. F. and Brodsky, J. L. (2014). The proteolytic landscape of the yeast vacuole. Cell Logist 4, e28023.

Hirano, H., Kimura, Y. and Kimura, A. (2015). Biological significance of co- and post-translational modifications of the yeast 26S proteasome. J Proteomics 134, 37– 46.

Hughes Hallett, J. E., Luo, X. and Capaldi, A. P. (2015). Snf1/AMPK promotes the formation of Kog1/Raptor-bodies to increase the activation threshold of TORC1 in budding yeast. eLife 4, e09181.

Hurley, J. H. (2015). ESCRTs are everywhere. EMBO J 34, 2398–2407.

Isasa, M., Katz, E. J., Kim, W., Yugo, V., González, S., Kirkpatrick, D. S., Thomson, T. M., Finley, D., Gygi, S. P. and Crosas, B. (2010). Monoubiquitination of RPN10 regulates substrate recruitment to the proteasome. Mol Cell 38, 733–745.

Katzmann, D. J., Stefan, C. J., Babst, M. and Emr, S. D. (2003). Vps27 recruits ESCRT machinery to endosomes during MVB sorting. J Cell Biol 162, 413–423.

Kim, J., Kundu, M., Viollet, B. and Guan, K.-L. (2011). AMPK and mTOR regulate autophagy through direct phosphorylation of Ulk1. Nat Cell Biol 13, 132.

Klionsky, D. J. Abdel-Aziz, A. K. Abdelfatah, S. Abdellatif, M. Abdoli, A. Abel, S. Abeliovich, H. Abildgaard, M. H. Abudu, Y. P. Acevedo-Arozena, A. et al. (2021). Guidelines for the use and interpretation of assays for monitoring autophagy (4th edition). Autophagy 17, 1–382.

Kors, S., Geijtenbeek, K., Reits, E. and Schipper-Krom, S. (2019). Regulation of proteasome activity by (post-)transcriptional mechanisms. Front Mol Biosci 6, 48.

Kushnirov, V. V. (2000). Rapid and reliable protein extraction from yeast. Yeast 16, 857–860.

Laporte, D., Salin, B., Daignan-Fornier, B. and Sagot, I. (2008). Reversible cytoplasmic localization of the proteasome in quiescent yeast cells. J Cell Biol 181, 737–745.

Lauwers, E., Jacob, C. and André, B. (2009). K63-linked ubiquitin chains as a specific signal for protein sorting into the multivesicular body pathway. J Cell Biol 185, 493–502.

Lecker, S. H., Goldberg, A. L. and Mitch, W. E. (2006). Protein degradation by the ubiquitin–proteasome pathway in normal and disease states. J Am Soc Nephrol 17, 1807–1819.

Li, J., Breker, M., Graham, M., Schuldiner, M. and Hochstrasser, M. (2019). AMPK regulates ESCRT-dependent microautophagy of proteasomes concomitant with proteasome storage granule assembly during glucose starvation. PLoS Genet 15, e1008387.

Li, J., Fuchs, S., Zhang, J., Wellford, S., Schuldiner, M. and Wang, X. (2016). An unrecognized function for COPII components in recruiting the viral replication protein BMV 1a to the perinuclear ER. J Cell Sci 129, 3597–3608.

Li, J. and Hochstrasser, M. (2020). Microautophagy regulates proteasome homeostasis. Curr Genet 66, 683–687.

Li, Y., Tomko, R. J. and Hochstrasser, M. (2015). Proteasomes: isolation and activity assays. Current protocols in cell biology / editorial board, Juan S. Bonifacino … [et al.] 67, 3.43.1-3.43.20.

MacDonald, C., Shields, S. B., Williams, C. A., Winistorfer, S. and Piper, R. C. (2020). A cycle of ubiquitination regulates adaptor function of the Nedd4-family ubiquitin ligase Rsp5. Curr Biol 30, 465–479.e5.

Marshall, R. S., McLoughlin, F. and Vierstra, R. D. (2016). Autophagic turnover of inactive 26S proteasomes in yeast is directed by the ubiquitin receptor Cue5 and the Hsp42 chaperone. Cell Rep 16, 1717–1732.

Marshall, R. S. and Vierstra, R. D. (2018). Proteasome storage granules protect proteasomes from autophagic degradation upon carbon starvation. eLife 7, e34532.

Mruk, D. D. and Cheng, C. Y. (2011). Enhanced chemiluminescence (ECL) for routine immunoblotting: An inexpensive alternative to commercially available kits. Spermatogenesis 1, 121–122.

Mumberg, D., Müller, R. and Funk, M. (1995). Yeast vectors for the controlled expression of heterologous proteins in different genetic backgrounds. Gene 156, 119–122.

Narayanaswamy, R., Levy, M., Tsechansky, M., Stovall, G. M., O’Connell, J. D., Mirrielees, J., Ellington, A. D. and Marcotte, E. M. (2009). Widespread reorganization of metabolic enzymes into reversible assemblies upon nutrient starvation. Proc Natl Acad Sci USA 106, 10147–10152.

Nemec, A. A., Howell, L. A., Peterson, A. K., Murray, M. A. and Tomko, R. J. (2017). Autophagic clearance of proteasomes in yeast requires the conserved sorting nexin Snx4. J Biol Chem 292, 21466–21480.

Oku, M., Maeda, Y., Kagohashi, Y., Kondo, T., Yamada, M., Fujimoto, T. and Sakai, Y. (2017). Evidence for ESCRT- and clathrin-dependent microautophagy. J Cell Biol 216, 3263–3274.

Papa, F. R. and Hochstrasser, M. (1993). The yeast DOA4 gene encodes a deubiquitinating enzyme related to a product of the human tre-2 oncogene. Nature 366, 313–319.

Piper, R. C., Dikic, I. and Lukacs, G. L. (2014). Ubiquitin-dependent sorting in endocytosis. Cold Spring Harb Perspect Biol 6, a016808.

Ren, X. and Hurley, J. H. (2010). VHS domains of ESCRT-0 cooperate in high-avidity binding to polyubiquitinated cargo. EMBO J 29, 1045–1054.

Sagot, I. and Laporte, D. (2019). The cell biology of quiescent yeast – a diversity of individual scenarios. J Cell Sci 132, jcs213025.

Schmidt, O. and Teis, D. (2012). The ESCRT machinery. Curr Biol 22, R116–R120.

Schmitt, S. M., Neslund-Dudas, C., Shen, M., Cui, C., Mitra, B. and Dou, Q. P. (2016). Involvement of ALAD-20S proteasome complexes in ubiquitination and acetylation of proteasomal α2 subunits. J Cell Biochem 117, 144–151.

Segev, N. (2020). ESCRTing proteasomes to the lysosome. PLoS Genet 16, e1008631.

Shah, S. S. and Kumar, S. (2021). Adaptors as the regulators of HECT ubiquitin ligases. Cell Death Differ 28, 455–472.

Shields, S. B., Oestreich, A. J., Winistorfer, S., Nguyen, D., Payne, J. A., Katzmann, D. J. and Piper, R. (2009). ESCRT ubiquitin-binding domains function cooperatively during MVB cargo sorting. J Cell Biol 185, 213–224.

Shields, S. B. and Piper, R. C. (2011). How ubiquitin functions with ESCRTs. Traffic 12, 1306–1317.

Sikorski, R. S. and Hieter, P. (1989). A system of shuttle vectors and yeast host strains designed for efficient manipulation of DNA in Saccharomyces cerevisiae. Genetics 122, 19–27.

Starita, L. M., Lo, R. S., Eng, J. K., von Haller, P. D. and Fields, S. (2012). Sites of ubiquitin attachment in Saccharomyces cerevisiae. Proteomics 12, 236–240.

Swaminathan, S., Amerik, A. Y. and Hochstrasser, M. (1999). The Doa4 deubiquitinating enzyme is required for ubiquitin homeostasis in yeast. Mol Biol Cell 10, 2583–2594.

Swaney, D. L., Beltrao, P., Starita, L., Guo, A., Rush, J., Fields, S., Krogan, N. J. and Villén, J. (2013). Global analysis of phosphorylation and ubiquitylation cross-talk in protein degradation. Nat Methods 10, 676–682.

Thibaudeau, T. A. and Smith, D. M. (2019). A practical review of proteasome pharmacology. Pharmacol Rev 71, 170–197.

Tokuyasu, K. T. (1973). A technique for ultracryotomy of cell suspensions and tissues. J Cell Bio 57, 551–565.

Tomko Jr, Robert J. and Hochstrasser, M. (2014). The intrinsically disordered Sem1 protein functions as a molecular tether during proteasome lid biogenesis. Mol Cell 53, 433–443.

Tomko, R. J. and Hochstrasser, M. (2013). Molecular architecture and assembly of the eukaryotic proteasome. Annu Rev Biochem 82, 415–445.

Velichutina, I., Connerly, P. L., Arendt, C. S., Li, X. and Hochstrasser, M. (2004). Plasticity in eucaryotic 20S proteasome ring assembly revealed by a subunit deletion in yeast. EMBO J 23, 500–510.

Vietri, M., Radulovic, M. and Stenmark, H. (2020). The many functions of ESCRTs. Nat Rev Mol Cell Biol 21, 25–42.

Waite, K. A., Mota-Peynado, A. D.-L., Vontz, G. and Roelofs, J. (2016). Starvation induces proteasome autophagy with different pathways for core and regulatory particles. J Biol Chem 291, 3239–3253.

Wang, G., Yang, J. and Huibregtse, J. M. (1999). Functional domains of the Rsp5 ubiquitin-protein ligase. Mol Cell Biol 19, 342–352.

Zhou, F., Wu, Z., Zhao, M., Murtazina, R., Cai, J., Zhang, A., Li, R., Sun, D., Li, W., Zhao, L. et al. (2019). Rab5-dependent autophagosome closure by ESCRT. J Cell Biol 218, 1908–1927.

## Supporting References

Amerik, A., Sindhi, N. and Hochstrasser, M. (2006). A conserved late endosome–targeting signal required for Doa4 deubiquitylating enzyme function. J Cell Biol 175, 825–835.

Chen, P., Johnson, P., Sommer, T., Jentsch, S. and Hochstrasser, M. (1993). Multiple ubiquitin-conjugating enzymes participate in the in vivo degradation of the yeast MATα2 repressor. Cell 74, 357–369.

## Supporting references

Ellison, M. J. and Hochstrasser, M. (1991). Epitope-tagged ubiquitin. A new probe for analyzing ubiquitin function. J Biol Chem 266, 21150–7.

Hodgins, R. R., Ellison, K. S. and Ellison, M. J. (1992). Expression of a ubiquitin derivative that conjugates to protein irreversibly produces phenotypes consistent with a ubiquitin deficiency. J Biol Chem 267, 8807–8812.

